# Condensation of PRPP amidotransferase by phase separation promotes *de novo* purine synthesis through intermolecular intermediate channeling in yeast

**DOI:** 10.1101/2023.06.05.543815

**Authors:** Masak Takaine, Rikuri Morita, Yuto Yoshinari, Takashi Nishimura

## Abstract

*De novo* purine synthesis (DPS) is up-regulated under conditions of high purine demand to ensure the production of genetic materials and chemical energy, thereby supporting cell proliferation. However, the regulatory mechanisms governing DPS remain largely unclear. We herein show that PRPP amidotransferase (PPAT), the rate-limiting enzyme in DPS, forms dynamic and motile condensates in *Saccharomyces cerevisiae* cells under a purine-depleted environment. The formation of condensates requires phase separation, which is driven by target of rapamycin complex 1 (TORC1)-induced ribosome biosynthesis. The self-assembly of PPAT molecules facilitates condensate formation, with intracellular PRPP and purine nucleotides both regulating this self-assembly. Moreover, molecular dynamics simulations suggest that clustering-mediated PPAT activation occurs through intermolecular substrate channeling. Cells unable to form PPAT condensates exhibit growth defects, highlighting the physiological importance of condensation. These results suggest that PPAT condensation is an adaptive mechanism that regulates DPS in response to both TORC1 activity and cellular purine demands.

**Highlights:**

- PRPP amidotransferase (PPAT) in budding yeast forms dynamic intracellular condensates in response to environmental purine deprivation.
- Ribosome crowding in the cytoplasm, induced by TORC1, drives the assembly of PPAT condensates.
- The purified PPAT protein alone condensates into submicron-sized particles *in vitro* under molecular crowding conditions.
- Purine nucleotides inhibit the self-assembly of PPAT, while PRPP promotes this process and counteracts the inhibition.
- The condensation of PPAT facilitates the intermolecular channeling of intermediates, thereby enhancing *de novo* purine synthesis.

## Introduction

Purine nucleotides and their derivatives play crucial roles in many cellular processes, making them essential metabolites for all living organisms. The purine nucleotide triphosphates, ATP and GTP, act as cellular energy currencies. In addition, purine metabolites are integrated into genetic materials, such as DNA and RNA, and into coenzymes, including NAD and coenzyme A. Purine nucleotides are synthesized through either salvage or *de novo* pathways. Under normal conditions, cells primarily synthesize the majority of purine nucleotides via the salvage pathway by adding phosphoribosyl pyrophosphate (PRPP), derived from the pentose phosphate pathway, to purine bases, which are often imported from outside the cell. Since at least one of the two pathways is required for purine synthesis, they provide a fail-safe mechanism for each other.

The *de novo* purine biosynthetic pathway consists of ten highly conserved sequential chemical reactions facilitated by six enzymes that convert PRPP to inosine 5′-monophosphate (IMP). IMP is then converted into adenosine 5′-monophosphate (AMP) or guanine 5′-monophosphate (GMP) through two branched paths. Among these enzymes, PRPP amidotransferase (PPAT) (EC 2.4.2.14) is the rate-limiting enzyme and plays a pivotal role in the regulation of *de novo* purine synthesis. PPAT catalyzes the initial step of the *de novo* pathway, namely, the transfer of amino groups from glutamine to PRPP to produce 5-phosphoribosylamine (PRA) (eq. 1). This reaction is further divided into two half-reactions: the release of ammonia (NH_3_) from glutamine (eq. 2) and the ligation of NH_3_ with PRPP (eq. 3). PPAT is a bifunctional enzyme with two catalytic domains, possessing an N-terminal glutamine amidotransferase (GATase) domain that catalyzes the former reaction, and a C-terminal phosphoribosyltransferase (PRTase) domain for the latter.

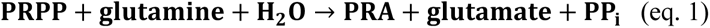

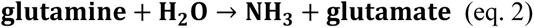

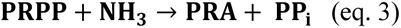

While the biochemical properties of each enzyme involved in purine synthesis have been extensively examined, the regulation of *de novo* purine synthesis within cells remains unclear. In bacterial and mammalian cells, when extracellular purine bases are available, purine nucleotides are preferentially synthesized by recycling bases through the salvage pathway, while the *de novo* pathway is strongly down-regulated^1^ ^2^. Conversely, under conditions of purine deprivation or high purine demand (*e.g.*, in actively proliferating cells), the *de novo* pathway is activated to provide sufficient levels of purine nucleotides^3^. Although the mechanisms underlying these regulatory processes have not yet been elucidated in detail, cells appear to limit the use of the *de novo* pathway in order to conserve energy and resources because the generation of each IMP requires up to five ATP and four amino acids.

The enzyme activity of PPAT is subject to feedback inhibition by the end products of the *de novo* pathway, such as AMP and GMP ^4–6^. This inhibition is considered to play a role in both suppressing *de novo* purine synthesis in the presence of salvageable extracellular purine bases and activating it during purine starvation ^4^. However, millimolar levels of AMP or GMP are required to inhibit PPAT catalytic activity^4, 6, 7^, whereas intracellular concentrations of these nucleotides are markedly lower ^8^, raising questions about the physiological relevance of this feedback inhibition. Other regulatory layers of PPAT activity, such as post-translational modifications and intracellular signaling, may also play a role, but have yet to be investigated.

Emerging evidence suggests that many metabolic enzymes, including PPAT, form condensates in response to external stimuli in order to regulate cellular metabolism ^9^. For example, the pyruvate kinase Cdc19 in budding yeast reversibly aggregates into large foci upon glucose starvation and heat shock ^10, 11^. This aggregation protects Cdc19 from stress-induced degradation and inactivates it, both of which are crucial for the efficient formation of stress granules and cell regrowth after stress. Similarly, PPAT reversibly localizes to cytoplasmic condensates upon the depletion of extracellular purine bases in budding yeast and HeLa cells ^12, 13^. Recent studies indicated that in HeLa cells, PPAT also colocalized with other enzymes involved in *de novo* purine synthesis within a multi-enzyme cellular complex known as a purinosome, which appeared to enhance pathway flux near mitochondria ^14^. However, the mechanisms regulating the formation of PPAT condensates and its physiological significance remain unclear.

We herein demonstrate that PPAT forms dynamic intracellular condensates through phase separation in budding yeast cells in the absence of external purine bases. Active ribosome synthesis promoted by target of rapamycin complex 1 (TORC1) is crucial for the molecular crowding that drives the phase transition. An *in vitro* analysis showed that the self-assembly of PPAT alone is sufficient for condensate formation, which is inhibited by purine nucleotides. Moreover, increased intracellular PRPP promotes the condensation of PPAT, opposing the inhibitory effects of purine nucleotides. Molecular dynamics (MD) simulations suggest the condensation-dependent activation of the enzyme by intermolecular NH_3_ channeling. Consistent with these results, PPAT condensates play a vital role in *de novo* purine synthesis to ensure cell proliferation. Collectively, we propose that PPAT condensation by phase transition is a rapid regulatory mechanism for *de novo* purine synthesis in response to both TORC1 activity and available purine bases.

## Results

### The yeast PPAT Ade4 concentrates in fine particles

Previous semi-comprehensive analyses revealed that the budding yeast PPAT Ade4 formed cytoplasmic foci in response to environmental purine bases^12, 15^. However, the dynamics of the assembly and disassembly of Ade4 foci, their physical properties, and the mechanisms underlying the formation of foci remain unclear. Therefore, we reexamined Ade4 foci using a cell biological approach. In the present study, adenine was used as a representative purine base because of its high solubility in medium and its role as a precursor for both AMP and GMP through reverse conversion from AMP to IMP (Fig. 1A). We visualized the localization of the Ade4 protein by tagging the chromosomal *ADE4* gene with a yeast codon-optimized GFP (hereafter referred to as GFP). As demonstrated in previous studies ^12, 15^, Ade4-GFP showed a large focus in cells transferred into medium lacking purine bases, but not in the presence of adenine (Fig. 1B). Unexpectedly, Ade4 tagged with a GFP bearing the monomeric mutation A206K^16^ (Ade4-mGFP) appeared as numerous fine dim dots under the same conditions (Fig. 1B). Consistently, Ade4 tagged with monomeric NeonGreen (Ade4-mNG) formed fine particles with markedly higher fluorescence intensities (Fig. 1B-C). Ade4 tagged with the monomeric red fluorescent protein mKate2 (Ade4-mKate2) showed similar particle localization (Fig. S1A). Therefore, we conclude that Ade4-GFP concentrates into 1-2 foci due to GFP dimerization.

**Figure 1.**
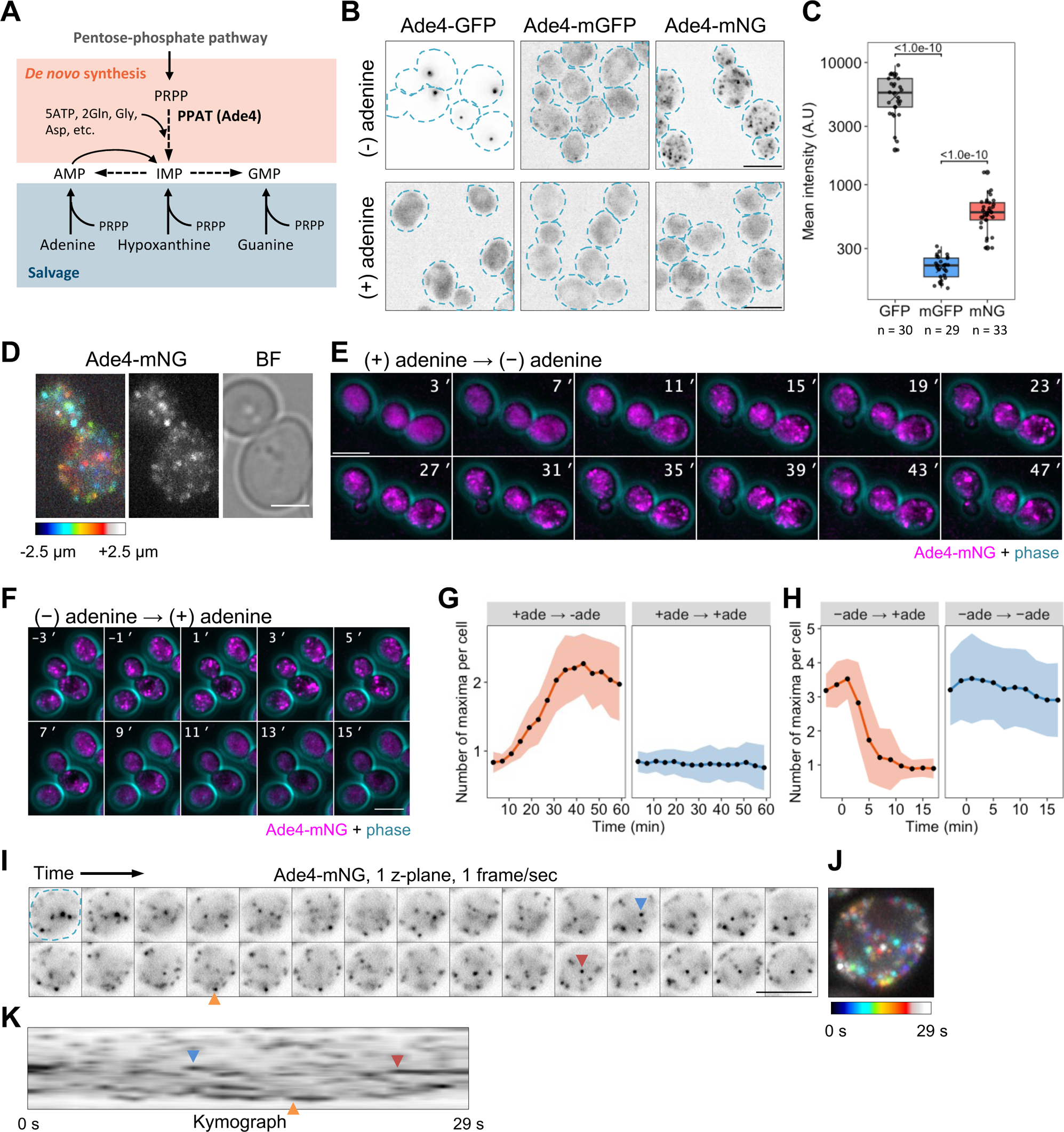
The yeast PPAT Ade4 concentrates in fine particles. A. Schematic of purine *de novo* and salvage pathways. B. Fluorescent images of cells expressing Ade4 tagged C-terminally with the indicated green fluorescent protein. Cells were grown in medium containing 0.02 mg/ml adenine, washed with medium with or without adenine, and then incubated in the same medium for 45 min before imaging. The intensity of GFP is represented by an inverted grayscale. Dotted lines indicate the cell outline. C. Quantification of the mean fluorescence intensity of Ade4 condensates observed in cells as shown in (B). Data are represented by box and dot plots. Cross marks indicate the population mean. *P*-values versus mGFP are shown (Steel’s multiple comparison test). D. A higher resolution image of Ade4-mNG particles. Z-positions are represented by a color code (left). A grayscale image of particles (middle) and a bright field image of cells (right) are also shown. E. Time-lapse imaging of the assembly of Ade4-mNG particles by adenine depletion. Cells were grown in medium containing 0.02 mg/ml adenine and washed with medium lacking adenine at t = 0 min. F. Time-lapse imaging of the disassembly of Ade4-mNG particles by adenine supplementation. Cells were grown in the absence of adenine and 0.02 mg/ml adenine was supplemented at t = 0 min. G, H. Panels G and H show the quantification of data in (E) and (F), respectively. The number of intracellular Ade4-mNG particles was counted as the number of maxima of green fluorescence intensity per cell. Data are the mean of 10-12 independent fields of view, and approximately 70 cells were examined per field. The shaded area indicates ± 1SD. I-K. Closer time-lapse imaging of Ade4-mNG particles. A cell bearing the particles (outlined by a dotted line in the first frame) was imaged on a single z-plane at 1-s intervals. The intensity of mNG is represented by an inverted grayscale. Panel J shows a color-coded superimposition of the images from each frame shown in (I). Panel K is a kymograph of the images shown in (I). Arrowheads indicate the appearance of some particles. Scale bars = 5 µm.

Z-stack imaging revealed that Ade4-mNG particles localized throughout the cytoplasm (Fig. 1D). Time-lapse imaging further demonstrated that these particles were dynamic structures: they assembled within 40 min after the depletion of extracellular adenine (Figs. 1E, G and S1B, and Movies S1-S3) and disassembled within 10 min after adenine supplementation (Figs. 1F, H and S1C, and Movies S4-S6). Closer time-lapse imaging indicated the high motility of particles: they moved freely in three dimensions and repeatedly moved in and out of the focal plane within seconds (Figs. 1I-K and S1D-F, and Movie S7). These results suggest that Ade4 forms dynamic and motile particles in response to the availability of purines.

In subsequent experiments, we primarily focused on Ade4-mNG particles to elucidate the molecular mechanisms regulating Ade4 condensation. We also examined the assembly of Ade4-GFP foci due to their similar properties to Ade4-mNG particles and their suitability for high-content imaging because of their higher brightness.

### Ade4 particles have liquid-like properties

The reversible assembly and dynamic nature of Ade4-mNG particles prompted us to hypothesize that they are liquid-like assemblies rather than static, solid-like aggregates. Therefore, we treated cells containing Ade4-mNG with 1,6-hexanediol, an aliphatic alcohol that dissolves various intracellular membrane-less structures ^17^. The permeabilization of yeast cells with digitonin did not affect the assembly of Ade4-mNG particles, whereas particles exposed to 10% 1,6-hexanediol plus digitonin significantly disassembled within 10 min, confirming their liquid-like properties (Fig. 2A-B).

**Figure 2.**
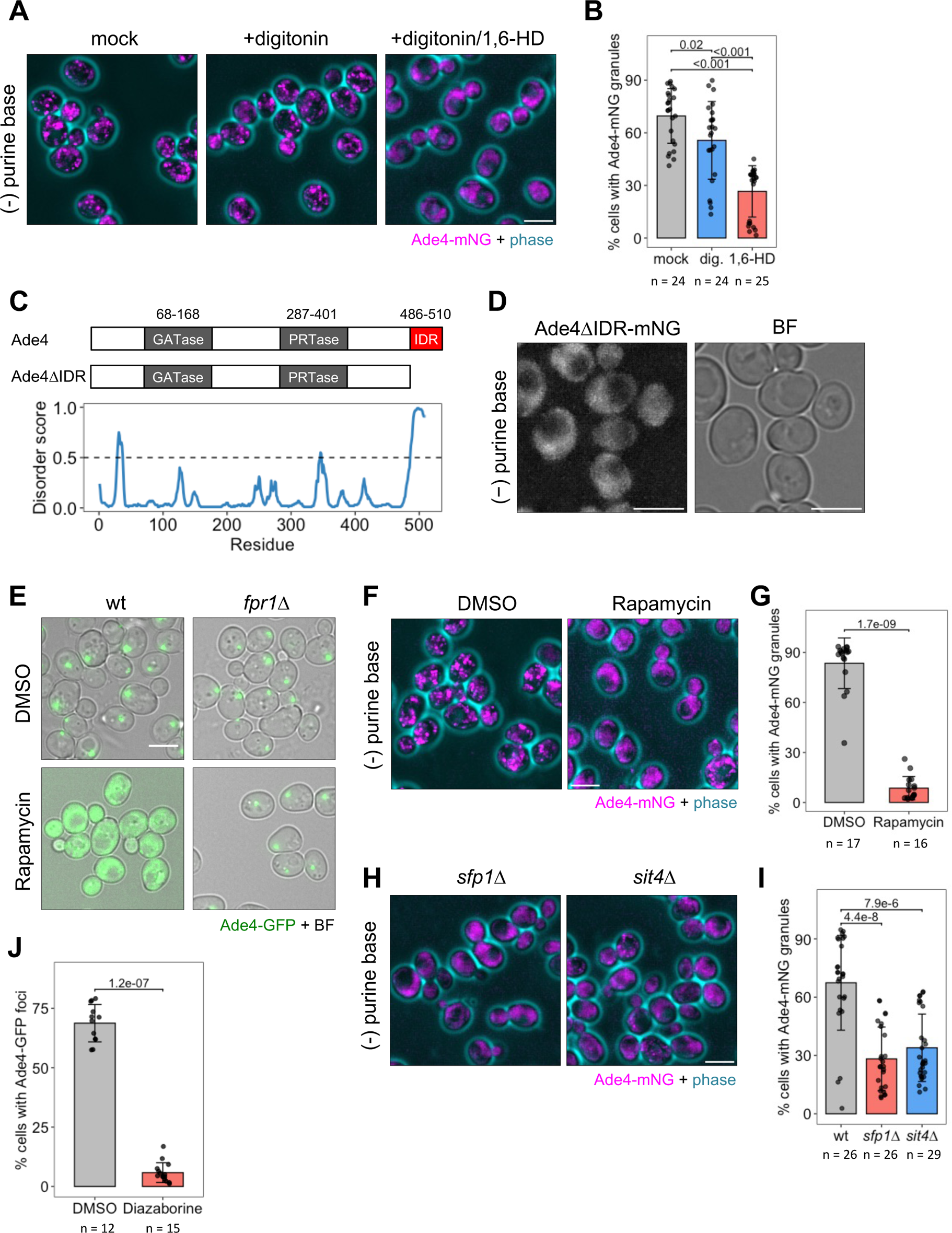
Ade4 condensate formation requires active TORC1 and ribosome biogenesis. A. The aliphatic alcohol 1,6–hexanediol dissolves Ade4 condensates. Wild-type cells harboring Ade4-mNG condensates were treated with 2 µg/ml digitonin ± 10% (w/v) 1,6- hexanediol for 10 min and then imaged. B. Quantification of data shown in (A). The percentage of cells with Ade4-mNG condensates per field of view (containing 361 ± 159 cells) was plotted. Data were pooled from three independent experiments. Bars and error bars indicate the mean ± 1SD. *P*-values were calculated using the two-tailed Steel-Dwass multiple comparison test. C. Domain structures of the budding yeast Ade4 and the IDR mutant. The bottom graph shows disorder predictions. D. The C-terminal IDR of Ade4 is required for condensate formation. Cells expressing the mutant Ade4 lacking the C-terminal IDR tagged with mNG were incubated in the absence of purine bases for 45 min and imaged. E. Wild-type and *fpr1*Δ cells expressing Ade4-GFP were grown in medium without adenine in the presence of 0.02% (v/v) DMSO or 0.02% DMSO plus 2 µg/ml rapamycin for 45 min before imaging. F. Rapamycin suppresses the assembly of Ade4-mNG condensates. Wild-type cells expressing Ade4-mNG were treated with rapamycin and imaged as described in (E). G. Quantification of data shown in (F). The percentage of cells with Ade4-mNG condensates per field of view (containing 443 ± 138 cells) was plotted. Data were pooled from two independent experiments. Bars and error bars indicate the mean ± 1SD. *P*-values were calculated using the two-tailed Mann-Whitney U test. H. The assembly of Ade4-mNG condensates was significantly suppressed in *sfp1*Δ and *sit4*Δ cells. I. Quantification of data shown in (H). The percentage of cells with Ade4-mNG condensates per field of view (containing 517 ± 123 cells) was plotted. Data were pooled from three independent experiments. Bars and error bars indicate the mean ± 1SD. *P*-values versus wt were calculated using the two-tailed Steel’s multiple comparison test. J. The inhibition of ribosome synthesis suppresses the assembly of Ade4 condensates. Cells expressing Ade4-GFP were grown in medium containing adenine and then incubated in medium without adenine in the presence of 0.05% (v/v) DMSO or 0.05% DMSO plus 5 µg/ml diazaborine for 45 min before imaging. The percentage of cells with Ade4-GFP foci per field of view (containing 780 ± 214 cells) was plotted. Data were pooled from two independent experiments. Bars and error bars indicate the mean ± 1SD. *P*-values were calculated using the two-tailed Mann-Whitney U test. Representative cell images are shown in Figure S3C. Scale bars = 5 µm.

Previous studies highlighted the role of an intrinsically disordered region (IDR) in the formation of intracellular membrane-less structures^18^. We analyzed the amino acid sequences of Ade4 and other PPATs using the DISOPRED algorithm^19^, a neural network-based method that predicts IDRs. We identified an IDR at the C terminus of Ade4 (Fig. 2C). The presence of the IDR was highly conserved across Ade4 homologues in Ascomycota and all fungi (Fig. S2A), as well as in PPATs from other kingdoms, including bacteria and mammals (Fig. S2B). However, the amino acid sequences of IDRs varied across species, even among yeasts (Fig. S2C-D). An Ade4 mutant lacking the IDR, Ade4ΔIDR-mNG, was incapable of forming punctate structures, suggesting the necessity of the IDR for Ade4 particle assembly (Fig. 2D). Collectively, these results suggest that Ade4 molecules condensate into fine particles through IDR-dependent phase separation.

### Formation of Ade4 condensates requires active TORC1 and ribosome biogenesis

We examined the factors required for phase separation, which facilitates the assembly of Ade4 condensates. Recent studies suggested that TORC1 controlled phase separation in the cytoplasm of both yeast and mammalian cells by modulating ribosome crowding^20^. Therefore, we investigated the role of TORC1 in the formation of Ade4 condensates. In cells, rapamycin binds to FKBP12, forming a complex that effectively inhibits TORC1^21^. The addition of rapamycin suppressed the assembly of Ade4-GFP foci in wild-type cells, but not in cells lacking Fpr1 (budding yeast FKBP1) (Fig. 2E). Similarly, rapamycin significantly inhibited the assembly of Ade4-mNG condensates in wild-type cells (Fig. 2F-G). These results indicate that TORC1 activity is essential for the formation of Ade4 condensates.

To further delineate the TORC1-related factors responsible for Ade4 condensate formation, we examined Ade4-GFP foci in 37 single-gene deletion mutants, each lacking one of the 17 upstream regulators, the 17 downstream effectors of TORC1 signaling, or 3 genes of interest (Fig. S3A-B). Consistent with the inhibitory effects of rapamycin, the assembly of Ade4-GFP foci was typically reduced in mutants of the upstream activators of TORC1 signaling (such as the Pib2 or EGO complex). In contrast, most of the downstream effectors of TORC1 signaling exerted negligible or no effects on the formation of Ade4-GFP foci. However, the formation of Ade4-GFP foci and Ade4-mNG condensates was significantly suppressed in *sfp1*Δ and *sit4*Δ cells (Figs. 2H-I and S3A-B).

Sfp1 is a transcriptional activator that regulates the expression of ribosomal protein and ribosome biogenesis genes ^22^. Sit4 is a subunit of type 2A protein phosphatase, the activity of which is controlled by TORC1 ^23^. Both have been implicated in cytoplasmic crowding through ribosome biogenesis downstream of TORC1 ^20, 24, 25^. Therefore, we evaluated the importance of ribosome synthesis on Ade4 condensate formation. Diazaborine specifically inhibits the maturation of the 60S ribosomal subunit and reduces the level of the 60S subunit in yeast cells ^26^. Treatment with diazaborine strongly suppressed the formation of Ade4-GFP foci and Ade4-mNG condensates, indicating that active synthesis of the ribosomal 60S subunit was required for the assembly of these condensates (Figs. 2J and S3C-D). Collectively, these results suggest that macromolecular crowding of the cytoplasm, driven by TORC1-mediated ribosome synthesis, is a primary mechanism facilitating the phase separation required for Ade4 condensation.

### Ade4 protein condensates into particles under molecular crowding conditions *in vitro*

Two hypotheses have been proposed for the molecular mechanisms regulating the assembly of Ade4 condensates. The first hypothesis suggests that Ade4 alone is sufficient for condensate formation, while the second posits that additional cellular components, such as proteins, lipids, and organelles, are required for Ade4 to condensate into particles. To distinguish between these possibilities, we investigated the properties of the purified Ade4 protein *in vitro*. Previous attempts to express Ade4 in its active form in bacteria were unsuccessful ^27^, prompting us to purify Ade4-mNG directly from budding yeast cells (Fig. S4A). Based on the findings of 20 quantitative proteomic studies, the intracellular concentrations of Ade4 were estimated to be approximately 0.5 µM (Fig. S5). In our *in vitro* assays, we selected concentrations of Ade4-mNG that were close to or slightly lower than this estimated value.

Under physiological ionic conditions (150 mM NaCl and 4 mM MgCl_2_, pH 7.5), purified Ade4-mNG dispersed homogenously, showing no signs of condensate formation (“0%” in Fig. 3A). We then examined the effects of macromolecular crowding using polyethylene glycol (PEG) as a crowding agent. The addition of PEG8000 led to the formation of sub-micron-sized particles of Ade4-mNG in a concentration-dependent manner; significant amounts of particles formed at concentrations higher than 10% PEG (Fig. 3A-C). Controls with the mNG protein alone did not form particles in 10% PEG (Fig. S4B). We also investigated the effects of different sizes of PEG and found that Ade4-mNG self-assembled into particles in the presence of PEG molecular weights higher than 4 kDa (Fig. S4C-E). Consistent with this size dependency, other crowding agents, such as 2% lysozyme (MW: 14 kDa) and Ficoll-400 (MW: 400 kDa), also induced the assembly of Ade4-mNG particles (Fig. S4F). These results suggest that Ade4 self-assembles into sub-micron particles that are similar in size to intracellular condensates under macromolecular crowding conditions.

**Figure 3.**
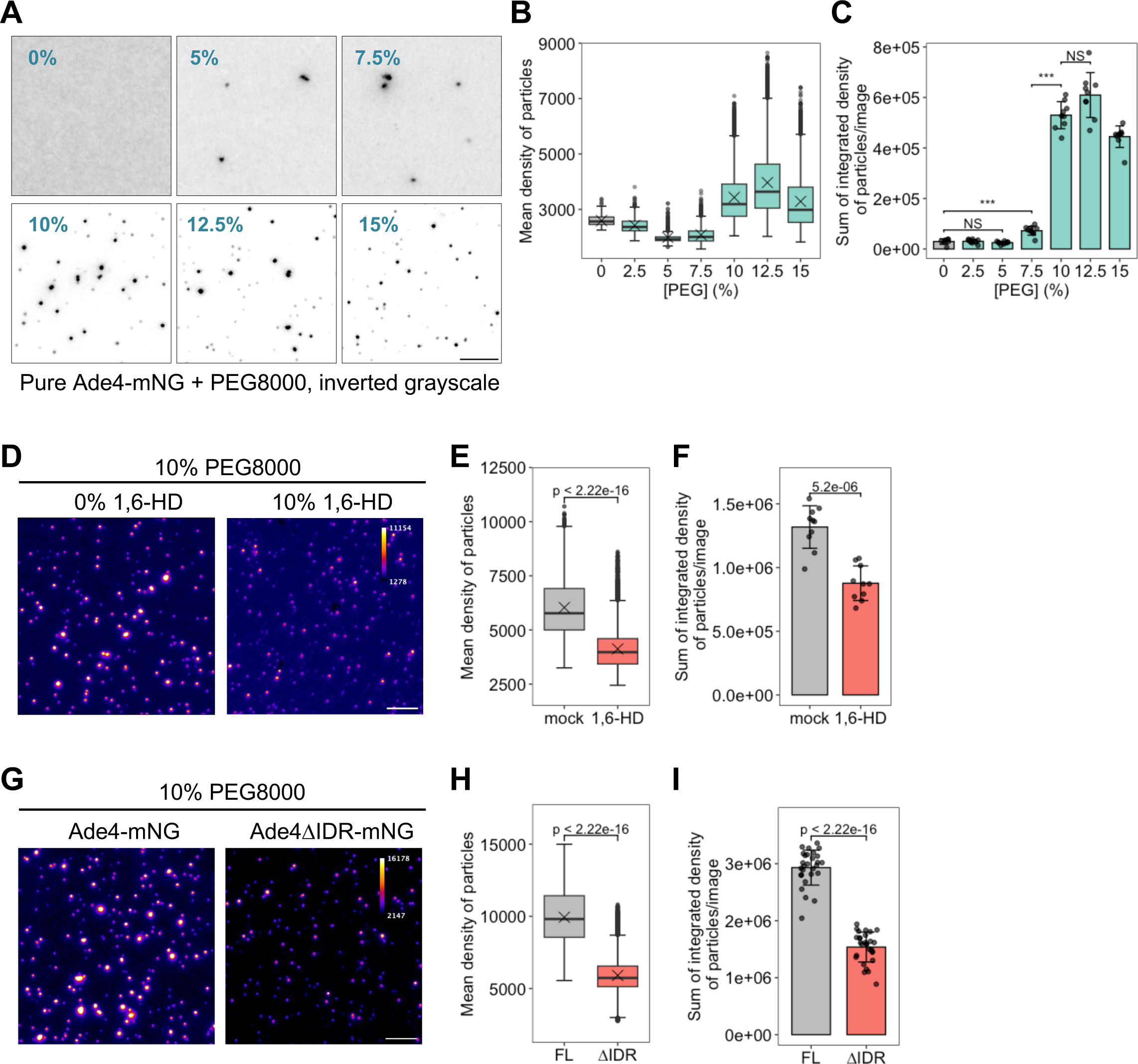
The Ade4 protein condensates into particles under molecular crowding conditions *in vitro*. A. Purified Ade4-mNG condensated into fine particles in a PEG concentration-dependent manner. Ade4-mNG (0.15 µM) was supplemented with PEG8000 at the indicated concentration (%, w/v). Inverted grayscale images are shown. B-C. Quantification of data shown in (A). The mean fluorescence intensities of Ade4-mNG particles were box plotted in (B). Cross marks indicate the population mean. The integrated fluorescence intensities of particles were summed per field and plotted in (C). Bars and error bars indicate the mean ± 1SD. Seven to 10 fields of view were imaged for each condition and an average of 390 particles per field were examined. *P*-values were calculated using the two-tailed Steel-Dwass multiple comparison test. D. 1,6-Hexanediol attenuated the *in vitro* condensation of Ade4-mNG. Ade4-mNG (0.15 µM) was supplemented with 10% PEG8000 in the presence of 0 or 10% (w/v) 1,6- hexanediol. Fluorescence intensity is represented by pseudo-colors. E-F. Quantification of data shown in (D). The mean fluorescence intensities of particles were box plotted in (E). Cross marks indicate the population mean. The integrated fluorescence intensities of particles were summed per field and plotted in (F). Bars and error bars indicate the mean ± 1SD. Ten fields of view were imaged for each condition, and an average of 1197 particles per field were examined. *P*-values were calculated using the two-tailed Mann-Whitney U test (E) or Welch’s *t*-test (F). G. Ade4ΔIDR-mNG condensated into particles less effectively than Ade4-mNG *in vitro*. The supplementation of 0.2 µM Ade4-mNG or Ade4ΔIDR-mNG with 10% PEG8000 was performed. Fluorescence intensity is represented by pseudo-colors. H-I. Quantification of data shown in (G). The mean fluorescence intensities of particles were box plotted in (H). Cross marks indicate the population mean. The integrated fluorescence intensities of particles were summed per field and plotted in (I). Bars and error bars indicate the mean ± 1SD. Twenty-eight fields of view were imaged for each condition, and an average of 1341 particles per field were examined. *P*-values were calculated using the two-tailed Mann-Whitney U test. Scale bars = 5 µm.

We also investigated the role of phase separation in the self-assembly of Ade4-mNG *in vitro*. In the presence of 1,6-hexanediol, the mean fluorescence intensity and total mass of Ade4-mNG particles were both markedly reduced (Fig. 3D-F). Consistently, the deletion of IDR from Ade4 markedly reduced condensate formation; however, the Ade4ΔIDR-mNG protein still formed visible particles (Fig. 3G-I). These results suggest that while Ade4 molecules form condensates independent of the C-terminal IDR moiety, phase separation is responsible for the complete assembly of Ade4 particles *in vitro*.

### Metabolomics analysis of budding yeast cells grown in the absence and presence of adenine

We investigated the mechanisms by which intracellular Ade4 condensates assemble in response to environmental purine deprivation. Nutrients such as amino acids and glucose typically regulate the activity of TORC1 ^28^. However, environmental purine bases, including adenine, do not affect TORC1 activity in mammalian cells ^29, 30^. Based on these findings, TORC1-mediated macromolecular crowding, which drives Ade4 condensate assembly, appears to be independent of extracellular purine base levels. Therefore, there may be additional layers of regulation that either suppress or promote condensate formation based on the availability of environmental purines. We hypothesized that metabolites related to the incorporated purine bases regulate the assembly of Ade4 condensates.

To identify these functional metabolites, we conducted a comprehensive metabolomics analysis using liquid chromatography-tandem mass spectrometry. Wild-type cells were cultured in the absence or presence of adenine for 1, 7, and >24 h and were then collected for metabolic profiling (Fig. 4A). A supervised classification method, a sparse partial least squares discriminant analysis (PLS-DA), demonstrated that samples were distinctly separated based on the presence of environmental adenine and the duration of exposure to adenine (Fig. 4B and Supplementary Data S1). A one-way analysis of variance (ANOVA) identified 56 metabolites whose concentrations were significantly affected by adenine (Fig. 4C and Supplementary Data S2). The metabolic profile changed over the duration of adenine exposure, indicating both short-and long-term responses (Fig. 4B-C). Intracellular concentrations of adenine increased more than five-fold within 1 h of the addition of adenine (Fig. 4D and Supplementary Data S3). A correlation analysis revealed that the concentrations of several metabolites, such as hypoxanthine, inosine, and deoxynucleotides, positively correlated with the change in intracellular adenine (Fig. 4E). Among them, hypoxanthine, which is directly converted from adenine, showed the greatest increase after adenine supplementation, and remained elevated (Fig. 4D). Meanwhile, purine nucleotides and their derivatives slightly increased in the presence of adenine, with only some of these changes being significant.

**Figure 4.**
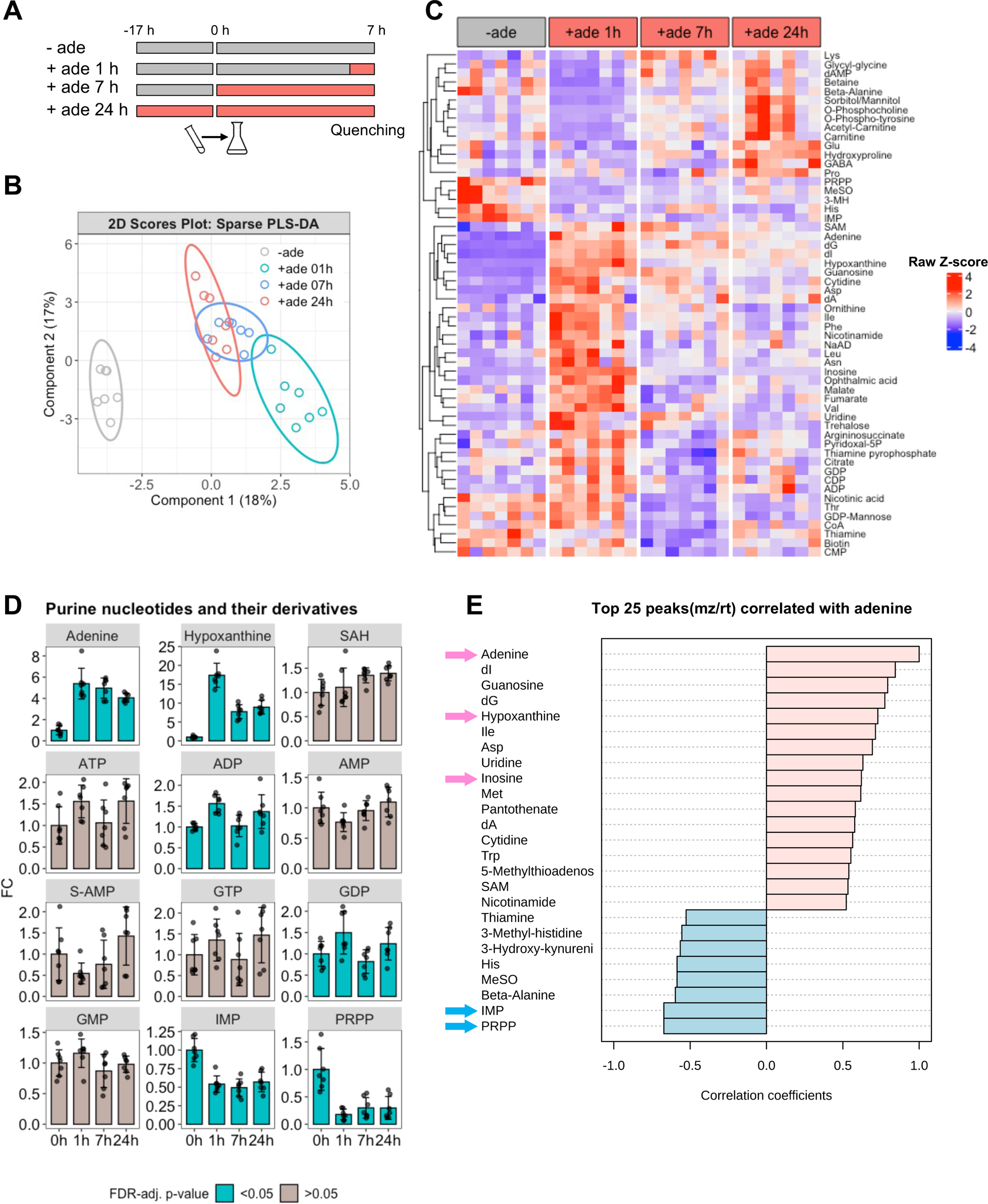
Metabolomics analysis of budding yeast cells grown in the absence and presence of adenine. A. Experimental scheme for adenine exposure and cell quenching. The preculture was inoculated into the main culture at 0 h. B. Sparse PLS-DA of metabolome samples. A 2D score plot of the first two components with ellipse plots (95% confidence level). C. Clustered heatmap of significantly changed metabolites. Rows indicate 56 metabolites that changed significantly between the indicated conditions (FDR-adjusted *p*-value <0.05). Columns represent seven independent samples for each condition. Values for each metabolite were converted to Z-scores and plotted. Metabolites were clustered using the complete linkage method and Euclidean distance measure. D. Changes in purine nucleotides and their derivatives. Data on the indicated metabolites were converted to fold change (FC) relative to the mean at 0 h (in the absence of adenine) and plotted. Bars and error bars indicate the mean ± 1SD calculated from 7 independent biological replicates for each condition. The color of the bar indicates whether the change in the metabolite was significant. E. Correlation coefficients of the top 25 metabolites correlated with intracellular adenine. Some metabolites of interest are indicated with arrows.

### Intracellular derivatives of purine nucleotides regulate the formation of Ade4 condensates

Since intracellular concentrations of adenine and hypoxanthine rapidly and significantly increased upon the addition of adenine, we speculated that these metabolites may suppress the formation of intracellular Ade4 condensates. However, these purine bases were converted to various metabolites after active uptake into cells through the purine-cytosine permease Fcy2^31^ (Fig. 5A). Therefore, we used mutant strains to clarify whether adenine and hypoxanthine suppressed Ade4 condensate formation. When wild-type cells were treated with medium containing either adenine or hypoxanthine, Ade4-GFP foci were completely disassembled (Fig. 5B-E, wt), indicating that hypoxanthine acted as an adenine substitute. In *aah1Δapt1Δ* cells, in which incorporated adenine was not metabolized further, the addition of adenine failed to disassemble Ade4 foci, whereas that of hypoxanthine completely disassembled foci (Fig. 5B-C), suggesting that adenine itself did not affect Ade4 condensates. Conversely, in *hpt1*Δ cells, in which incorporated hypoxanthine was not metabolized further, the addition of hypoxanthine failed to disassemble Ade4 foci (Fig. 5B, D), indicating that hypoxanthine itself also did not affect Ade4 condensates. Notably, adenine failed to completely disassemble Ade4-GFP foci in *hpt1*Δ cells, suggesting that adenine affected wild-type cells primarily through its metabolic conversion to hypoxanthine.

**Figure 5.**
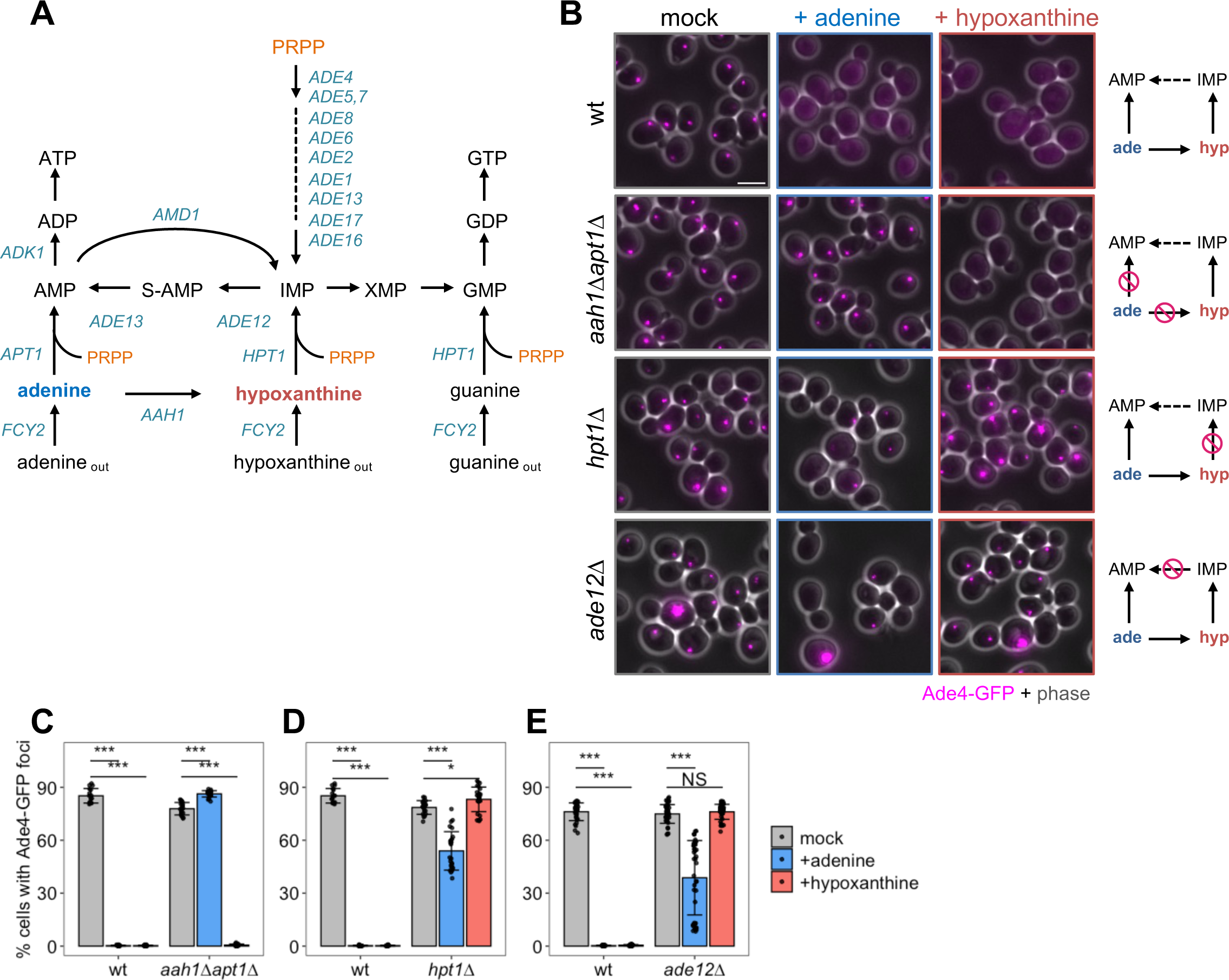
Intracellular derivatives of purine nucleotides regulate the formation of Ade4 condensates. A. *De novo* and salvage pathways for purine synthesis in budding yeast. B. Assembly of Ade4-GFP foci in purine metabolism mutants. Cells with Ade4-GFP foci in the absence of purine bases (mock) were treated with the same medium or medium containing 20 µg/ml adenine (+adenine) or 25 µg/ml hypoxanthine (+hypoxanthine) for 20 min and then imaged. Simplified views of the metabolism of the incorporated purine bases in the strains are shown. C-E. Quantification of data shown in (B). The percentage of cells with foci per field of view was quantified and plotted. Bars and error bars indicate the mean ± 1SD. Data were pooled from 2-3 independent experiments, and more than 16 fields of view were imaged for each condition. *P*-values versus the mock were calculated using the two-tailed Steel’s multiple comparison test: *, *p* <0.01; ***, *p* <10^-10^. Scale bar = 5 µm.

We also investigated the effects of purine nucleotide metabolism on condensate disassembly. In *ade12*Δ cells, in which IMP was not converted to AMP via adenylosuccinate (S-AMP), the addition of hypoxanthine failed to disassemble Ade4 foci, whereas adenine moderately decreased foci (Fig. 5B, E). The inhibition of Ade12 by alanosine, a specific inhibitor of adenylosuccinate synthetase ^32^, produced similar results (Fig. S7A-C). Therefore, we conclude that the anabolism of external purine bases is required to regulate the assembly of Ade4 condensates.

### Purine nucleotides attenuate the assembly of Ade4 condensates *in vitro*

The results of our genetic analyses suggest that AMP and related purine nucleotide derivatives play a vital role as functional metabolites that regulate the formation of Ade4 condensates. To directly test this possibility, we examined the inhibitory effects of purine nucleotides on the *in vitro* assembly of Ade4-mNG condensates. In the presence of GMP, AMP, ADP, and ATP, the condensation of Ade4-mNG was significantly reduced (Fig. 6A-B and Fig. S8C). AMP, ADP, and ATP were more effective than GMP, while S-AMP had no significant effect. Concentrations higher than 250 µM of AMP, ADP, or ATP were required to achieve maximal inhibitory effects, whereas the inhibitory effects of GMP plateaued at 100 µM (Fig. 6C and Fig. S8D-F). Consistent with our *in vivo* results (Fig. 5), adenine did not affect the *in vitro* condensation of Ade4 (Fig. S8A-B). Similarly, the ribose-bound hypoxanthine derivatives, inosine and deoxyinosine (dI), had no impact on Ade4 condensation (Fig. S8A-B). S-adenosylhomocysteine (SAH), a derivative of S-adenosylmethionine, exerted approximately one-third the inhibitory effect of AMP (Fig. S8G-H). Importantly, AMP also inhibited the condensation of Ade4ΔIDR-mNG (Fig. 6D). These results suggest that purine nucleotides, including AMP, ADP, ATP, and GMP, suppress the condensation of Ade4 molecules by inhibiting IDR-independent self-assembly.

**Figure 6.**
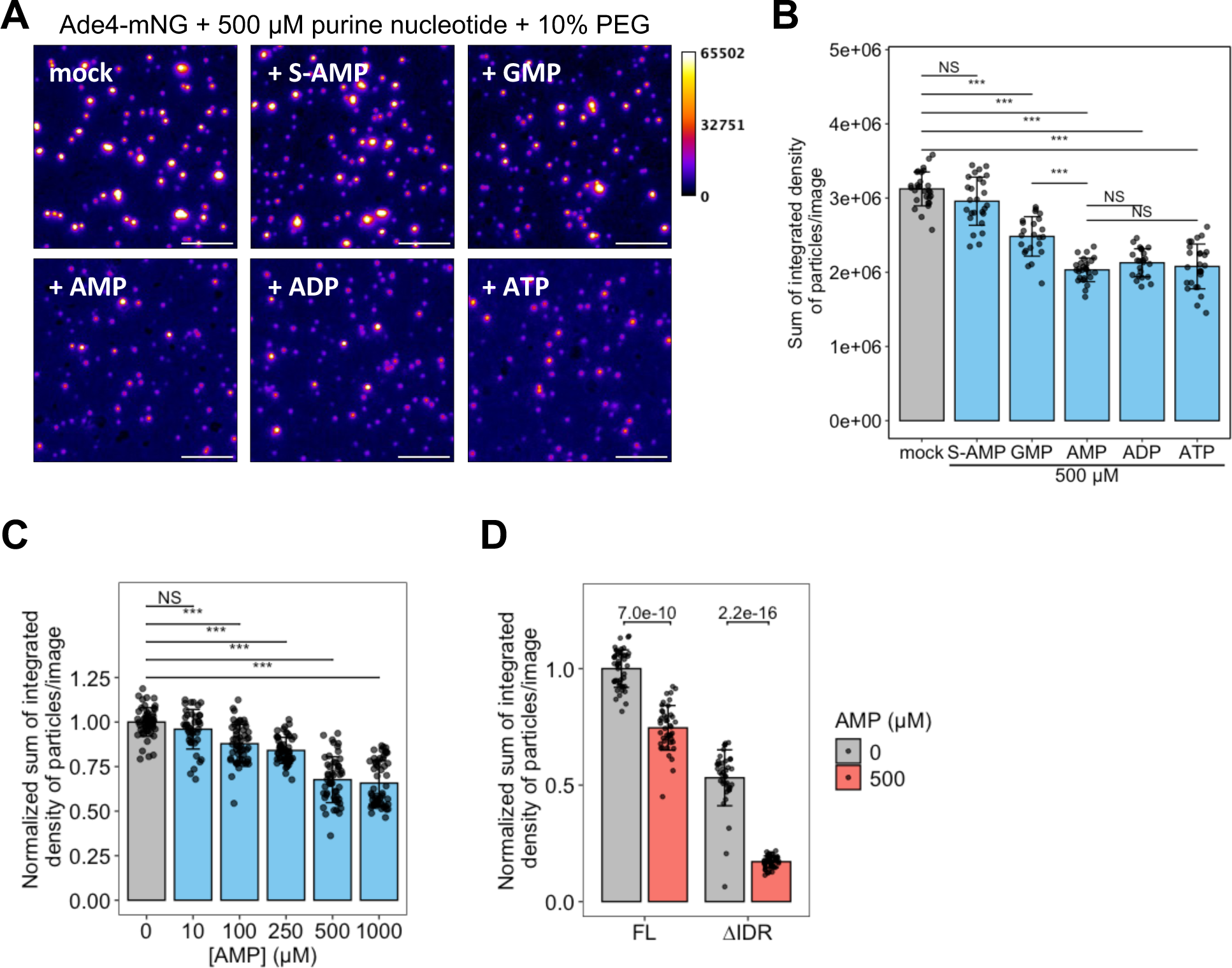
Purine nucleotides attenuate the assembly of Ade4 condensates *in vitro*. A. Effects of purine nucleotides on the *in vitro* assembly of Ade4 condensates. Ade4-mNG (0.2 µM) was supplemented with 10% PEG8000 in the presence of the indicated purine nucleotide. Fluorescence intensity is represented by pseudo-colors. Scale bars = 5 µm. B. Quantification of data shown in (A). The integrated fluorescence intensities of condensates were summed per field and plotted. Bars and error bars indicate the mean ± 1SD. Between 23 and 28 fields of view were imaged for each condition from two independent preparations, and an average of 1323 particles per field were examined. *P*-values were calculated using the two-tailed Tukey-Kramer’s multiple comparison test. C. The supplementation of 0.2 µM Ade4-mNG with 10% PEG8000 was performed in the presence of the indicated concentration of AMP. The integrated fluorescence intensities of the condensates were summed per field, normalized to the mean at 0 µM, and plotted. Bars and error bars indicate the mean ± 1SD. From 4 independent preparations, 41-55 fields of view were imaged for each condition, and an average of 1200 particles per field were examined. *P*-values were calculated using the two-tailed Steel’s multiple comparison test. D. Effects of AMP on the *in vitro* assembly of Ade4 and Ade4ΔIDR condensates. The supplementation of 0.2 µM Ade4-mNG (FL) or Ade4ΔIDR-mNG (ΔIDR) with 10% PEG8000 was performed in the presence of 0 or 500 µM AMP. The integrated fluorescence intensities of condensates were summed per field, normalized to the mean of FL at 0 µM AMP, and plotted. Bars and error bars indicate the mean ± 1SD. Between 40 and 45 fields of view were imaged for each condition from 3 independent preparations. *P*-values were calculated using the two-tailed Welch’s *t*-test.

### PRPP facilitates the assembly of Ade4 condensates *in vitro* and *in vivo*

The inhibitory effects of purine nucleotides on the *in vitro* formation of Ade4 condensates led us to hypothesize that these nucleotides may similarly regulate the formation of intracellular Ade4 condensates. However, the limited extent of these inhibitory effects suggested the existence of additional regulatory factors that promote the condensation of Ade4. The intracellular concentrations of these factors may decrease with adenine supplementation and increase when adenine is absent. Our correlation analysis identified several metabolites whose changes inversely correlated with intracellular adenine levels (Fig. 4E). Among them, we focused on IMP, which is directly synthesized from hypoxanthine, and PRPP, which directly interacts with Ade4 as a substrate (Fig. 4D). We found that IMP did not affect the assembly of Ade4 condensates *in vitro*, even at concentrations that were higher than those in yeast cells ^8^ (Fig. S9A-B). In contrast, PRPP concentrations higher than 50 µM significantly increased the mean fluorescence intensity and total mass of *in vitro* Ade4 condensates under 10% PEG conditions (Figs. 7A and S10A). Quantitative analyses revealed that adenine supplementation reduced intracellular PRPP levels from approximately 30 μM to less than 10 μM (Fig. S6B), guiding our selection of PRPP concentrations within the range of 10–200 μM for subsequent assays. Notably, the promoting effects of PRPP were more evident at lower PEG concentrations, at which Ade4 condensates rarely formed; the addition of 100 µM PRPP enabled the formation of Ade4 condensates in 5% and 7.5% PEG (Fig. 7B-C and S10B-C). These results suggest that PRPP promotes and augments the assembly of Ade4 condensates *in vitro*. Furthermore, the inhibitory effect of 1 mM AMP on Ade4 condensation was effectively counteracted by PRPP at concentrations higher than 50 µM, with this effect being more pronounced in 7.5% PEG than in 10% PEG (Fig. 7D, E and S10D), suggesting that PRPP antagonizes the inhibitory effect of AMP.

**Figure 7.**
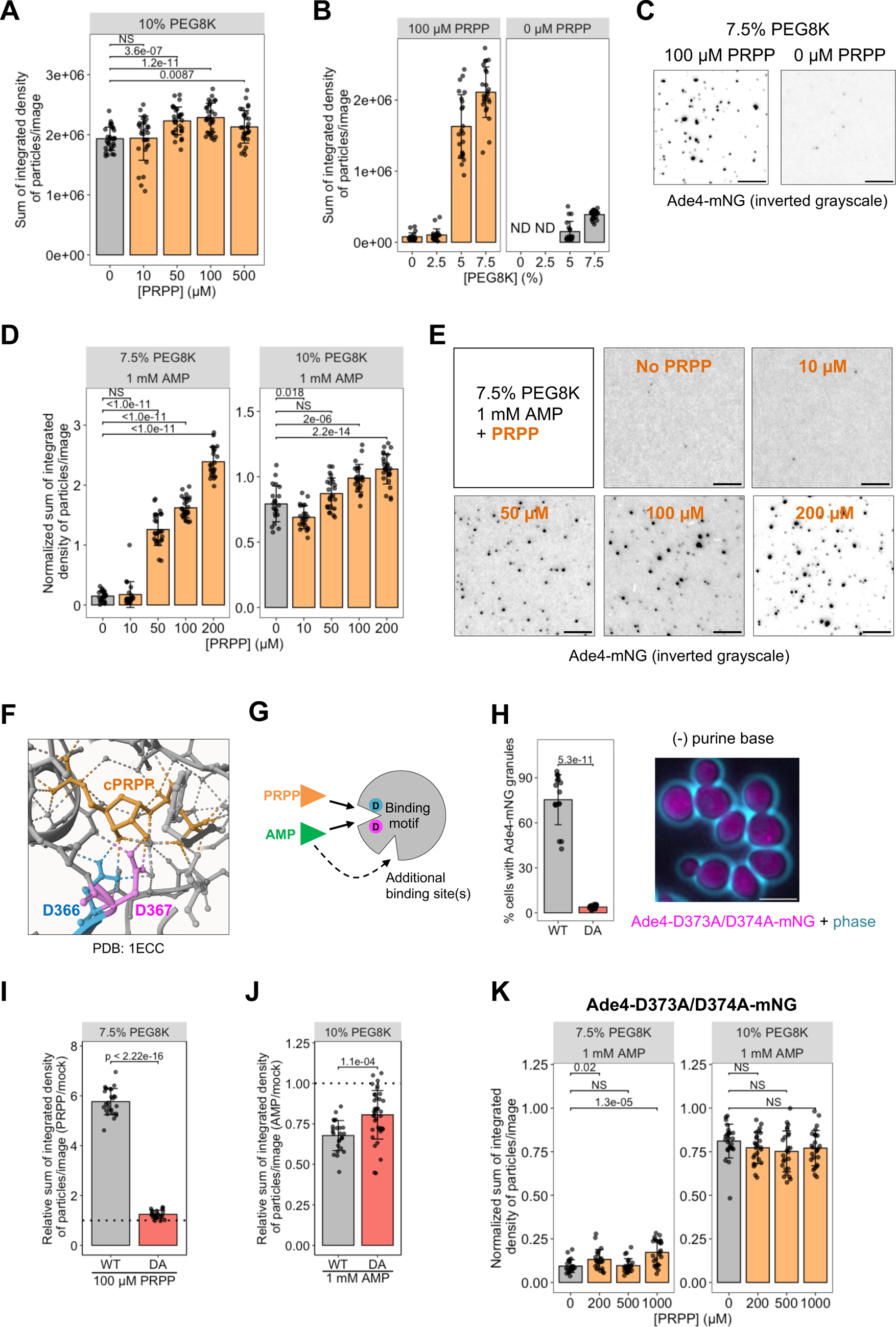
PRPP facilitates the assembly of Ade4 condensates. A. PRPP augments the *in vitro* assembly of Ade4 condensates. The supplementation of 0.2 µM Ade4-mNG with 10% PEG8000 was performed in the presence of the indicated concentration of PRPP. The integrated fluorescence intensities of the condensates were summed per field and plotted. Throughout the figure, bars and error bars indicate the mean ± 1SD. From 2 independent preparations, 29-31 fields of view were imaged for each condition, and an average of 1200 particles per field were examined. *P*-values were calculated using the two-tailed Steel’s multiple comparison test. B, C. PRPP promotes the formation of Ade4 condensates *in vitro* at suboptimal PEG concentrations. In the presence of 100 or 0 µM PRPP, 0.2 µM Ade4-mNG was supplemented with PEG8000 at the indicated concentration and imaged. The integrated fluorescence intensities of the condensates were summed per field and plotted in (B). From 2 independent preparations, 19-24 fields of view were imaged for each condition. ND, not determined. Representative images at 7.5% PEG are shown in (C). D, E. PRPP antagonizes the inhibitory effects of AMP. In the presence of 0 or 1 mM AMP and the indicated concentrations of PRPP, 0.2 µM Ade4-mNG was supplemented with 7.5% or 10% PEG8000. The integrated fluorescence intensities of the condensates were summed per field, normalized to the mean of the mock control (no AMP and no PRPP), and plotted in (D). From 2 independent preparations, 21-27 fields of view were imaged for each condition. *P*-values were calculated using the two-tailed Steel’s multiple comparison test. Representative images at 7.5% PEG are shown in (E). F. Interactions between a PRPP analog and specific amino acid residues in the PRPP binding motif in the *E. coli* PPAT (pdb file: 1ecc). The dotted line indicates a hydrogen bond. G. PRPP and purine nucleotides (*e.g.*, AMP) interact competitively with the binding motif. Purine nucleotides may also interact with additional binding site(s) and inhibit Ade4 condensation. See the text for details. H. The interaction of Ade4 with PRPP is important for condensate assembly. Ade4-mNG (WT) and Ade4-D373A/D374A-mNG (DA) cells were incubated in the absence of purine bases for 45 min and imaged. The percentages of cells with condensates per one field of view were quantified and plotted. Data were pooled from 2 independent experiments, and more than 16 fields of view were imaged for each condition. *P*-values were calculated using the two-tailed Mann-Whitney U test. A representative image of mutant cells is shown. I. PRPP did not stimulate the condensate assembly of the DA mutant. The supplementation of 0.2 µM Ade4-mNG (WT) and Ade4-D373A/D374A-mNG (DA) with 7.5% PEG8000 was performed in the presence of 0 or 100 µM PRPP and imaged. The integrated fluorescence intensities of the condensates were summed per field, normalized to the mean at 0 µM PRPP (dotted line), and plotted. From 2 independent preparations, >22 fields of view were imaged for each condition. *P*-values were calculated using the two-tailed Welch’s *t*-test. J. The DA mutant is less sensitive to the inhibitory effects of AMP. The supplementation of 0.2 µM Ade4-mNG (WT) and Ade4-D373A/D374A-mNG (DA) with 10% PEG8000 was performed in the presence of 0 or 1 mM AMP and imaged. The integrated fluorescence intensities of the condensates were summed per field, normalized to the mean at 0 mM AMP (dotted line), and plotted. From 2-3 independent preparations, >24 fields of view were imaged for each condition. *P*-values were calculated using the two-tailed Mann-Whitney U test. K. PRPP did not antagonize AMP in the condensate formation of the DA mutant. In the presence of 0 or 1 mM AMP and the indicated concentrations of PRPP, 0.2 µM Ade4-D373A/D374A-mNG was supplemented with 7.5% or 10% PEG8000. The integrated fluorescence intensities of the condensates were summed per field, normalized to the mean of the mock control (no AMP and no PRPP), and plotted. From 2 independent preparations, 22-25 fields of view were imaged for each condition. *P*-values were calculated using the two-tailed Steel’s multiple comparison test. Scale bars = 5 µm.

We then examined the effects of PRPP on the formation of cellular Ade4 condensates. The 13-residue PRPP binding motif at the catalytic site is well conserved among PPATs of various species, including budding yeast, with two adjacent aspartic acids that are crucial for PRPP binding ^33^ (Fig. S10E). A structural analysis of *E. coli* PPAT showed that these aspartic acids formed hydrogen bonds with carbocyclic PRPP (cPRPP), a stable analog of PRPP ^34^ (Fig. 7F). This motif also interacted with purine nucleotides in bacterial PPATs ^35–37^ (Figs. 7G and S10F), partially explaining the competitive inhibition of the enzyme by purine nucleotides. We introduced mutations at these aspartic acids (D373A and D374A, referred to as Ade4-DA) and investigated whether the interaction with PRPP affected the assembly of Ade4 condensates. Ade4-DA-mNG was virtually incapable of forming punctate structures (Fig. 7H), demonstrating that the interaction with PRPP was essential for the assembly of Ade4 condensates in cells.

Further purification of the mutant Ade4 protein allowed us to examine its *in vitro* biochemical properties (Fig. S10G). Ade4-DA-mNG condensated into particles under 10% PEG conditions, similar to wild-type Ade4-mNG (Fig. S10H), indicating that the mutation did not impair the self-assembly of Ade4 under crowding conditions. We found that PRPP only negligibly promoted the assembly of Ade4-DA-mNG condensates (Fig. 7I), suggesting that the binding of PRPP at the catalytic site was responsible for facilitating the condensation of Ade4 molecules. In contrast, the DA mutant remained sensitive to AMP; AMP inhibited the assembly of Ade4-DA-mNG condensates, albeit less effectively than that of Ade4-mNG condensates (Fig. 7J). Consistently, PRPP was unable to counteract the inhibitory effects of AMP on the condensation of Ade4-DA, even at concentrations higher than 500 μM (Figs. 7K and S10I-J). Therefore, while the interaction of purine nucleotides at the PRPP binding motif partially contributed to the inhibition of Ade4 condensation, purine nucleotides also appeared to suppress Ade4 condensation through additional binding sites (Fig. 7G).

### MD calculations predict the possible condensation-dependent activation of PPAT by enabling intermolecular substrate channeling

The induction of Ade4 condensate formation by the substrate PRPP prompted us to propose that the condensation of Ade4 promotes its enzymatic activity. In glutamine-dependent PRA synthesis (Gln-PRA activity, Fig. 8A), NH_3_ is initially released from glutamine at the GATase site and must then be transferred to PRPP at the PRTase site. A structural analysis of *E. coli* PPAT suggested that PRPP binding creates a transient hydrophobic channel linking the GATase and PRTase sites ^34^, thereby facilitating the efficient intramolecular transfer of NH_3_ between these catalytic sites (Figs. 8B-C and S11A). However, this intramolecular substrate channeling may be independent of the condensation state of the enzyme. In addition to glutamine-derived NH_3_, PPAT also directly utilize ambient NH_3_ as a nitrogen donor to produce PRA, referred to as NH_3_-dependent PRA synthesis (NH_3_-PRA activity, Figs. 8A and S11B) ^6^. Based on a condensation-dependent activation mechanism, we hypothesized that the Ade4 condensate may enable the intermolecular transfer of NH_3_ between molecules (Fig. 8C, right).

**Figure 8.**
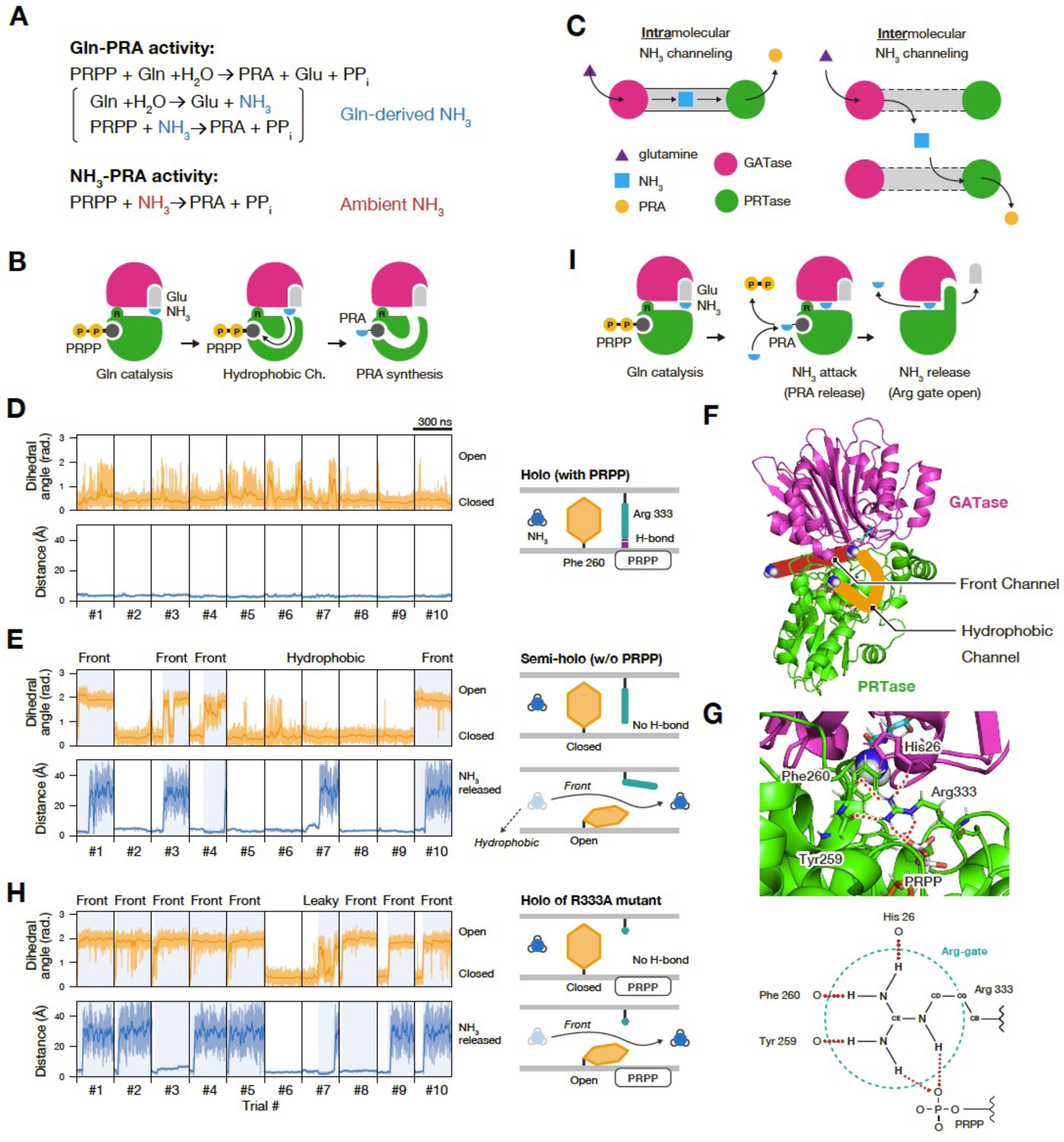
MD calculations predict the possible condensation-dependent activation of PPAT by enabling intermolecular substrate channeling. A. Reaction formulae corresponding to the glutamine-dependent PRA synthesis (Gln-PRA activity) and NH_3_-dependent PRA synthesis (NH_3_-PRA activity) of PPAT. B. Reaction mechanisms suggested in a previous study. C. Intramolecular and intermolecular channeling of glutamine-derived NH_3_. D, E, H. Time series for the dihedral angle of the Phe260 side chain and the distance of NH_3_ from the initial structure (D: holo; E: semi-holo; and H: R333A mutant). Moving averages are plotted in bold lines. Under each condition, 300-ns simulations were performed ten times individually. The blue backgrounds indicate the open state of the Phe260 sidechain. The schematic diagram shows the state of the Arg gate and the movement of the NH_3_ molecule. F. The hydrophobic (orange) and front (red) channels of *E. coli* PPAT. G. The conserved Arg residue functions as a gate of the front channel. Top: a structural snapshot around the Arg gate. Bottom: a scheme of the Arg gate. Hydrogen bonds are shown as red dotted lines. I. The reaction mechanisms suggested in the present study.

To test this hypothesis, we investigated the behavior of NH_3_ on the PPAT molecule by conducting MD simulations using the crystal structure of *E. coli* PPAT in its active form ^34^. MD simulations for the holo state of PPAT, which included glutamate, NH_3_, PRPP, and Mg^2+^, showed that NH_3_ mostly remained at the initial or adjacent positions during ten trials and did not reach the catalytic site, despite the presence of the hydrophobic channel (Fig. 8D). This result suggests a low frequency of intramolecular channeling relative to our simulation time. In contrast, in the semi-holo state containing only glutamate and NH_3_, NH_3_ escaped through the hydrophobic channel once in ten trials. Unexpectedly, NH_3_ was instead released from the molecule through a transient channel directed towards Arg333 in four out of ten trials (Fig. 8E-F and Movie S8). This Arg333 is conserved in yeast Ade4 and is referred to as the front channel. Prior to the release of NH_3_, we noted that the hydrophobic side chain of Phe260 flipped from its initial structure. Furthermore, Arg333 functioned as a gate for NH_3_ by forming hydrogen bonds with the backbone oxygen of Tyr259, Phe260, His26, and the phosphate group of PRPP (Fig. 8G). Notably, the absence of PRPP increased structural fluctuations in the Arg gate in the semi-holo state, thereby increasing the likelihood of gate opening.

Additional simulations on the Arg333Ala mutant under holo conditions confirmed the critical role of this Arg gate. The flipping of Phe260 and subsequent release of NH_3_ were observed in six out of ten trials on the Arg333Ala mutant, despite the presence of PRPP at the catalytic site (Fig. 8H). This increased gate-opening frequency in the mutant suggests that the normally closed Arg gate, coupled with PRPP, suppresses the flipping of the Phe260 side chain and the release of NH_3_. Since the PRTase catalytic site utilize ambient NH_3_ as a nitrogen donor ^6^, the release of PRA may simultaneously facilitate the release of internal glutamine-derived NH_3_ through the front channel (Fig. 8I and S11C). Collectively, these simulation results support PPAT condensation facilitating the intermolecular transfer of leaked NH_3_ between closely located molecules, leading to the efficient utilization of glutamine-derived NH_3_ (Figs. 8C, right panel and S11D).

### Formation of Ade4 condensates is important for efficient *de novo* purine synthesis during cell growth

Authentic Gln-PRA activity and NH_3_-PRA activity (Fig. 8A) have both been implicated in the *de novo* biosynthesis of purine nucleotides *in vivo* ^38^ ^39^. Since NH_3_-PRA activity depends on the concentration of NH_4_^+^ in medium, a fraction of which is present as NH_3_ ^6^, Ade4 condensates may be critical for Gln-PRA activity under low ammonium conditions. To examine this possibility, we compared the growth of cells expressing wild-type Ade4 with those expressing the Ade4ΔIDR mutant, which is unable to form condensates *in vivo*. On solid medium without adenine, the growth of the 3×HA-tagged Ade4ΔIDR cells was markedly slower than that of wild-type Ade4 cells (Fig. 9A). The protein level of Ade4ΔIDR grown in adenine-free medium was similar to that of wild-type Ade4 (Fig. 9B). Supplementation with adenine restored the growth of the *ADE4ΔIDR* mutant to wild-type rates, indicating that the slow growth of the mutant was due to reduced enzyme activity rather than dominant-negative inhibition by the mutant protein. Notably, the growth of the *ADE4ΔIDR* mutant significantly decreased with reductions in the concentration of ammonium ions from ammonium sulfate (AS) in the medium (Fig. 9A). We further quantified cell growth by measuring growth curves in liquid medium. The *ADE4ΔIDR* mutant exhibited significantly slower growth, particularly in the absence of ammonium ions, while the growth rate of the wild-type strain remained almost independent of the ammonium ion concentration in the medium (Fig. 9C-D). Collectively, these results suggest that the condensation of Ade4 into particles is critical for *de novo* purine synthesis, particularly under low ammonium conditions, where Gln-PRA activity plays a pivotal role.

**Figure 9.**
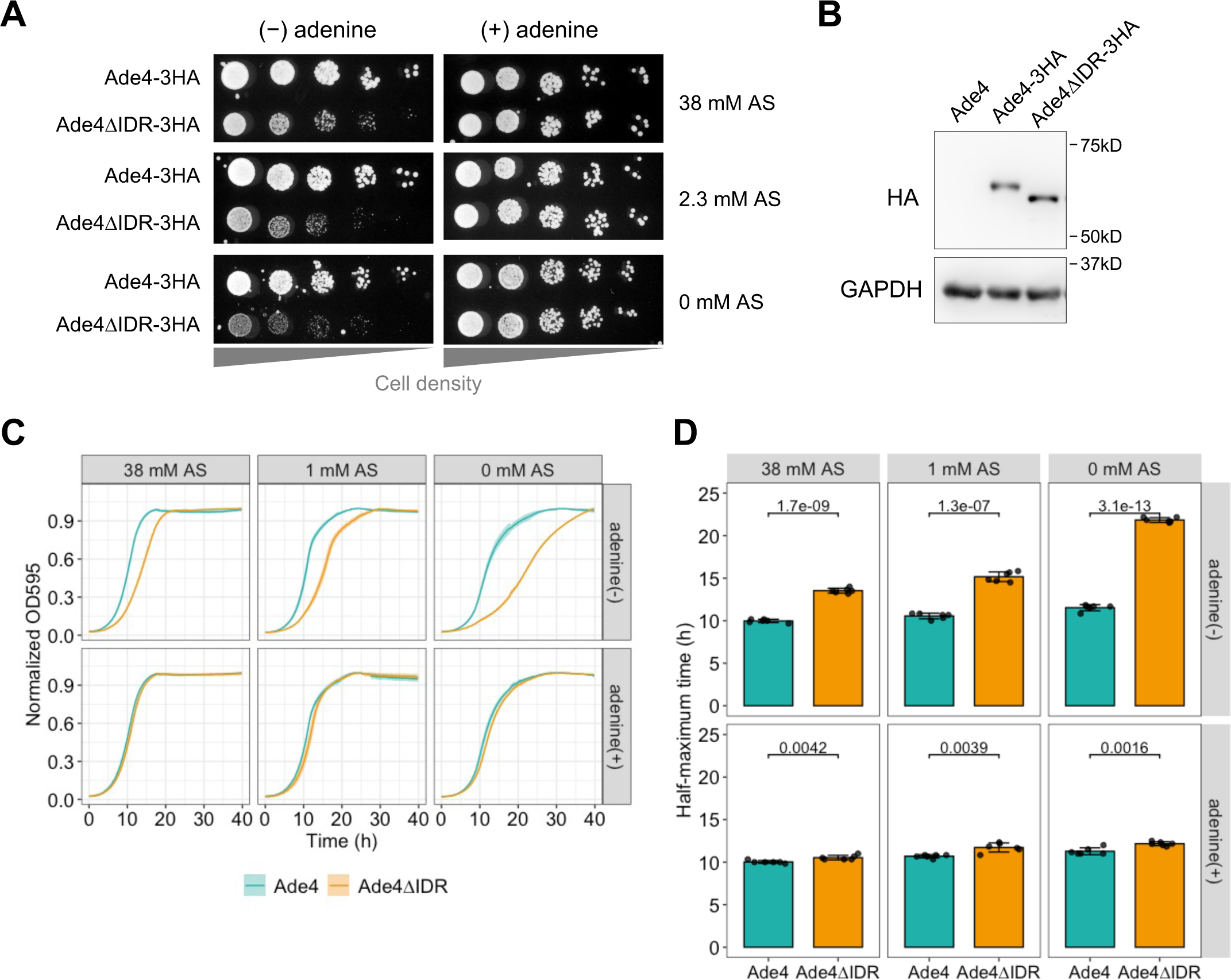
The formation of Ade4 condensates is important for efficient *de novo* purine synthesis during cell growth. A. Growth of ADE4-3HA and ADE4ΔIDR-3HA cells on solid media. Cells of each strain were spotted on medium containing the indicated concentration of ammonium sulfate (AS), and then grown at 30°C for 2-3 days. B. Immunoblot analysis of the protein level of Ade4. Total cell extracts were prepared from cells expressing the indicated Ade4 construct. Ade4-3HA and Ade4ΔIDR-3HA were detected as approximately 60-kDa bands by an anti-HA antibody. The bands of GAPDH are shown as a loading control. C. Growth curves of Ade4-3HA and Ade4ΔIDR-3HA cells in liquid media without adenine. Cells were grown at 30°C for 40 h in medium containing the indicated concentration of ammonium sulfate (AS) in the absence or presence of adenine. Data are the mean of 6 replicates, and the shaded area indicates ± 1SD. D. Quantification of data shown in (C). The time to half-maximum density (corresponding to normalized OD_595_ = 0.5) was calculated and plotted. Bars and error bars indicate the mean ± 1SD. *P*-values were calculated using the two-tailed Welch’s *t*-test.

## Discussion

We demonstrated that the budding yeast PPAT forms intracellular condensates through TORC1-mediated macromolecular crowding and PRPP-dependent self-assembly to regulate *de novo* purine synthesis in response to extracellular purine bases (Fig. 10). MD simulations revealed a previously unidentified pathway by which an intermediate NH_3_ derived from glutamine leaks from the PPAT molecule (Figs. 8I and S11C). We propose that Ade4 condensation enhances Gln-PRA activity by facilitating a domino-like chain reaction through intermolecular intermediate channeling. The condensation of PPAT, which has two catalytic domains, is considered to be equivalent to the co-clustering of two distinct enzymes, GATase and PRTase, in a confined space. Although NH_3_ is expected to diffuse rapidly throughout the cytoplasm ^40^, within Ade4 condensates—where the enzyme concentration is high—another enzyme may capture NH_3_ and effectively process it by the PRTase domain (Figs. 8C, right panel and S11D). This form of enzyme activation by intermolecular NH_3_ channeling corresponds to clustering-mediated metabolic channeling theoretically predicted by Castellana et al. ^41^. The present study is the first demonstration that a native enzyme may be activated by this type of metabolic channeling in eukaryotic cells.

**Figure 10.**
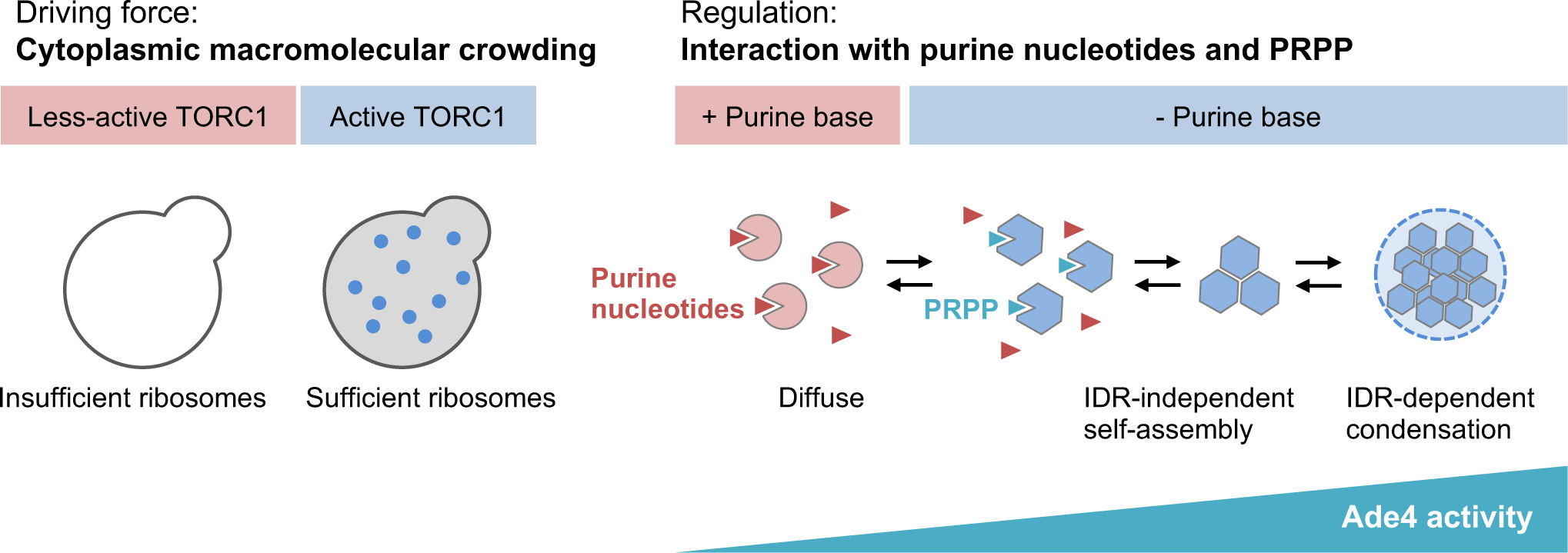
Molecular crowding and interactions with effectors regulate Ade4 condensation. A schematic model showing the regulation of Ade4 condensation proposed in the present study. See the text for details.

Recent studies revealed that many metabolic enzymes form intracellular condensates by phase separation, allowing cellular metabolism to adapt to variable growth conditions^9^. These condensates serve as temporary stores for inactive enzymes, helping cells survive adverse environmental conditions^9^. In some cases, filament formation by the enzyme enhances catalytic activity^42, 43^. Our results add another example where the condensation of a metabolic enzyme boosts catalytic activation through intermolecular substrate channeling. Notably, Ade4 condensates are unique among the cellular condensates reported to date due to their dynamic nature; they assemble and disassemble quickly and reversibly in response to environmental purine bases and exhibit high motility.

Ade4 (PPAT) condensates resemble enzyme clusters known as purinosomes, which form reversibly in HeLa cells under purine deprivation ^13^. Purinosomes enhance the channeling of metabolic intermediates in the *de novo* pathway near mitochondria, thereby increasing pathway flux ^14^. However, these two structures differ in several respects. Purinosomes contain all six enzymes involved in *de novo* purine synthesis with formylglycinamidine ribonucleotide synthase as the core, whereas cellular clusters of PPAT are smaller and only some colocalize with purinosomes^44^. Furthermore, purinosomes are static, assembling and disassembling in approximately 1.5 and 1 h, respectively, and are non-motile^13^. Therefore, we hypothesize that PPAT condensates are structurally distinct from purinosomes and may coexist inside or outside of them. Since PPAT is the rate-limiting enzyme in *de novo* purine synthesis, its condensates may provide a rapid and flexible response to fluctuations in environmental purine base levels, whereas purinosomes may represent a more constant adaptation to purine scarcity.

We also found that purified Ade4 self-assembled into fine particles under molecular crowding conditions. *In vitro* phase separation assays further demonstrated that Ade4 condensation involved both IDR-independent self-assembly and IDR-dependent condensation. In addition to the C-terminal IDR, the Ade4 protein has a short putative IDR at the N terminus. Consistently, plant and insect PPATs have an IDR at the N terminus, but not at the C terminus, suggesting that both ends of the protein may contribute to condensate formation by phase separation. Therefore, multiple IDRs may be involved in complete condensate formation. Nevertheless, it is intriguing to speculate that C-terminal IDR-independent self-assembly is essential for subsequent IDR-dependent condensation, where phase separation is required for stable condensation into particles (see Fig. 10, right panel).

The enzymatic activity of PPAT is considered to be regulated in a feedback manner by millimolar levels of AMP and GMP ^45^. However, intracellular purine mononucleotide concentrations are generally maintained in the micromolar range, from tens to hundreds, and are not markedly affected by extracellular purine bases (Fig. 4D and previous studies^46–48^). Therefore, this feedback regulation appears to be physiologically irrelevant, at least in response to environmental purine bases. In contrast, the present results indicate that purine nucleotides inhibit the IDR-independent self-assembly of Ade4, while PRPP promotes condensate formation *in vitro*. We speculate that these two regulators control the assembly of Ade4 condensates, thereby affecting PPAT enzyme activity *in vivo*. Since ADP and ATP, similar to AMP, suppressed the assembly of Ade4 condensates, GDP and GTP may also exert similar inhibitory effects to those of GMP. Therefore, the concentrations of all purine nucleotides may be physiologically relevant and sufficient to exert inhibitory effects on Ade4 condensate formation in cells.

The intracellular concentrations of purine nucleotides were previously shown to slightly increase in the presence of environmental purine bases ^46–48^, while that of PRPP decreased to approximately 10 μM. At this level, PRPP is unable to interact with Ade4 in order to promote its assembly, and, thus, purine nucleotides may predominantly interact with Ade4 and prevent condensate formation. In the absence of purine bases, PRPP levels may increase up to several tens of micromolar, thereby promoting the condensation of Ade4. Since PRPP-docked Ade4 is prone to condensation, the local concentration of PRPP within condensates is expected to markedly increase. Consistent with this notion, the *K*_m_ of PPAT for PRPP in yeast was previously reported to be 110 μM^49^, which is 3.7-fold higher than the intracellular concentration of PRPP even in the absence of adenine. Similarly, in mammals, intracellular PRPP concentrations range between 5 and 30 μM^50^, while the enzyme’s *K*_m_ for PRPP is 480 μM^51^. The enrichment of PRPP within Ade4 condensates provides an example of the spatial control of substrate concentrations by biomolecular condensates^52^.

Consistent with the present results, the binding of PRPP to the specific binding motif at the catalytic site of *E. coli* PPAT has been reported to induce a structural change in the enzyme ^34, 53^, which may contribute to the enhanced clustering of the enzyme under molecular crowding conditions. AMP also interacts with the PRPP binding motif; however, in contrast to PRPP, it inhibits self-assembly. Therefore, competitive binding to the motif explains the mechanism by which PRPP counteracts the effects of purine nucleotides. However, since the Ade4-DA mutant remains sensitive to inhibition by AMP, further structural analyses are required to elucidate the assembly mechanisms and effects of purine nucleotides.

It is important to note that the addition of purine bases to culture media was found to reduce the concentration of PRPP in both bacterial cells ^50^ and mammalian cells ^54^, suggesting a conserved mechanism across species. The most likely explanation for this is the consumption of PRPP, coupled with the conversion of incorporated purine bases by hypoxanthine-guanine phosphoribosyltransferase (HGPRT, known as Hpt1 in budding yeast). HGPRT utilizes PRPP as a substrate more efficiently than PPAT, as evidenced by its markedly lower *K*_m_ for PRPP^55^. Consistently, HGPRT-negative lymphoblasts from Lesch-Nyhan syndrome patients showed 4- to 8-fold higher intracellular PRPP levels than those from normal individuals^56^, which supports this notion and is consistent with the phenotype of *hpt1Δ* cells observed in the present study.

We found that ribosome biogenesis downstream of TORC1 signaling was required for Ade4 condensate formation. These results align with the theory that TORC1 regulates macromolecular crowding of the cytoplasm through ribosome synthesis, which, in turn, controls biomolecular condensation by phase separation ^20^. The present study is the first experimental demonstration of a metabolic enzyme assembling into condensates via the TORC1-regulated phase transition of the cytoplasm (see Fig. 10, left panel). Since TORC1 and PPAT IDRs are both conserved across many species, the condensation of PPAT by TORC1 may represent a common feature in eukaryotic cells.

The involvement of TORC1 is another crucial aspect of Ade4 condensates. TORC1 is activated by sensing nutrients, such as amino acids and glucose ^57^. Therefore, the requirement of TORC1 activity for PPAT condensation ensures that the *de novo* synthesis of purines is maximized under conditions that are favorable for cell growth. In mammals, TORC1 transcriptionally up-regulates the enzymes involved in the mitochondrial tetrahydrofolate cycle, thereby increasing *de novo* purine synthesis ^58^. Correlating the metabolic flux of purine synthesis with the nutritional status via TORC1 activity appears to be a conserved and multi-layered regulatory mechanism for cell proliferation.

Disorders in the regulation of purine metabolism are associated with various diseases, such as cancer, hyperuricemia (gout), and immunological defects ^59, 60^. A more detailed examination of purine metabolism will provide insights into the molecular mechanisms underlying these diseases. Importantly, a recent study implicated the up-regulation of PPAT in the malignant progression of many cancer types ^61^. Therefore, elucidating the regulatory mechanism of PPAT activity will not only enhance our understanding of purine metabolism in a biological context, but also contribute to advances in basic and therapeutic research on cancer.

## Materials and Methods

### Yeast strains and plasmids

The yeast strains, plasmids, and primers used in the present study are listed in Supplementary Tables S1, S2, and S3, respectively. These strains were constructed by a PCR-based method ^62^ and genetic crosses. The yeast knockout strain collection was originally purchased from GE Healthcare (cat. # YSC1053).

Regarding the construction of plasmids used for the C-terminal tagging of monomeric GFP or mKate2, DNA fragments encoding the corresponding fluorescent proteins were artificially synthesized (Eurofin Genomics, Ebersberg, Germany). These fragments were replaced with GFP of pYM25^62^ using *Sal*I/*Bgl*II sites to yield MTP3114 and MTP3131, respectively. The plasmid used for the C-terminal tagging of mNG-Strep tag II (MTP3139) was constructed in the same manner: a DNA fragment encoding single mNG fused with the Strep II tag (WSHPQFEK) by a spacer sequence (ELYKGSA) was synthesized and then replaced with the GFP of pYM25.

### Media and cell culture

Synthetic medium optimized for the growth of the BY strain was prepared as described by Hanscho et al. ^63^. Growth assays in solid and liquid media used optimized synthetic media lacking glutamate, phenylalanine, serine, and threonine. Yeast nitrogen base without amino acids, but with AS (cat. # Q30009) was purchased from Invitrogen. Yeast nitrogen base without amino acids and AS (cat. # CYN0501) was obtained from FORMEDIUM. Synthetic medium without ammonium sulphate was supplemented with 8 mM proline as an alternative nitrogen source. Rich medium was standard YPD medium composed of 1% (w/v) yeast extract (BD, cat. # 212750), 2% (w/v) bacto-peptone (BD, cat. # 211677), and 2% (w/v) D-glucose. Glucose (cat. # 049-31165), adenine sulfate (cat. # 018-10613), hypoxanthine (cat. # 086-03403), 1, 6-hexanediol (cat. # 087-00432), and digitonin (cat. # 043-2137) were purchased from FUJIFILM Wako. Cells were grown to the mid-log phase at 30°C before imaging unless otherwise noted. To replace medium containing adenine with medium lacking adenine, cells were collected in a microtube by brief centrifugation and suspended in medium lacking adenine. This washing procedure was repeated at least twice. Rapamycin was purchased from LC Laboratories (cat. # R- 5000).

### Microscopy

Cells expressing a GFP-fusion protein were concentrated by centrifugation and sandwiched between a slide and coverslip (No. 1.5 thickness, Matsunami, Osaka, Japan). Immobilized cells were imaged using an inverted fluorescent microscope (Eclipse Ti2-E, Nikon, Tokyo, Japan) equipped with a CFI Plan Apoλ 100× Oil DIC/NA1.45 objective lens and CMOS image sensor (DS-Qi2, Nikon). The fluorescent signal was collected from stacks of 11 *z*-sections spaced by 0.5 µm, and the maximum projections of optical sections were shown unless otherwise noted. A CFI Plan Apoλ 60× Oil Ph3 DM/NA1.40 or CFI Plan Apoλ 100× Oil Ph3 DM/NA1.45 objective lens was used for phase contrast imaging. Images of cells were acquired from several fields of view for each experimental condition. All digital images were processed with the Fiji image platform^64^.

### Quantitative analysis of intracellular fluorescent condensation

Intracellular fluorescent condensation (*i.e.*, Ade4-GFP foci or Ade4-mNG particles) was automatically detected and analyzed by running Fiji macros (https://github.com/masaktakaine/FP-foci-quantification and https://github.com/masaktakaine/FP-granules-quantification). Briefly, these macros extract cell outlines from phase-contrast images and detect fluorescent condensation inside the cell boundary using the *FindMaxima* function of Fiji software.

To monitor temporal changes in the number of intracellular Ade4-mNG particles by time-lapse imaging, the number of maxima of green fluorescence intensity in the field was counted in each time frame and divided by the total cell number by running a Fiji macro (https://github.com/masaktakaine/Counting-FP-granules-in-time-lapse). In cases in which phase-contrast images were unavailable, we manually examined condensation.

### Spot growth assay

Cells were grown to the mid-log phase at 30°C in synthetic medium, and cell density (cells/ml) was measured by an automated cell counter (TC20, Bio-Rad). The culture was diluted by medium lacking nitrogen and carbon sources to a density of 1×10^6^ cells/ml, serially diluted 5-fold, and spotted on plates. Plates were incubated at 30°C for 2-3 days and then imaged by an image scanner (GT-X830, EPSON).

### Measurement of growth curves in liquid cultures

Cell growth in liquid media was monitored in a 96-well flat-bottomed plate using a microplate reader (iMark, Bio-Rad). Cells were grown to the mid-log phase and then diluted to OD_600_ = 0.1. Each well was filled with 300 µl of the cell dilution and OD_595_ was measured at 30°C every 10 min.

### Immunoblot analysis

A denatured whole-cell extract was prepared according to the method reported by von der Haar (2007) ^65^. Briefly, mid-log cells were harvested from 2–3 ml of the culture by centrifugation. The cell pellet was resuspended in a small volume (<100 µl) of lysis buffer (0.1 M NaOH, 50 mM EDTA, 2% (w/v) SDS, and 2% (v/v) β-mercaptoethanol) and then incubated at 90°C for 10 min. The sample was neutralized by the addition of a one-tenth volume of 1 M acetic acid, incubated at 90°C for 10 min, and mixed with a one-third volume of 4×Laemmli protein sample buffer (cat. # 1610747, Bio-Rad). Following centrifugation at 20,000×*g* at 4°C for 10 min, the upper 90% of the supernatant was collected as the clarified whole-cell extract and stored at -30°C until used. Proteins were resolved by SDS-PAGE using a pre-cast gel containing 10% (w/v) acrylamide (SuperSep™ Ace, cat. # 195-14951, FUJIFILM Wako) and then transferred onto a PVDF membrane (Immobilon-P, Millipore) using a protein transfer system (Trans-Blot Turbo Transfer System, Bio-Rad). HA-tagged Ade4 proteins were detected using an anti-HA antibody (mouse monoclonal (12CA5), cat. # GTX16918, Funakoshi) at a 1:1000 dilution. GFP-tagged proteins were detected using an anti-GFP antibody (mouse monoclonal (1E10H7), cat. #66002-1-Ig, ProteinTech) at a 1:1000 dilution. GAPDH was detected using an anti-GAPDH antibody (mouse monoclonal (1E6D9), cat. # 60004-1-Ig, ProteinTech) at a 1:1000 dilution. PGK1 was detected using an anti-PGK1 antibody (mouse monoclonal (22C5D8), cat. # ab113687, Abcam) at a 1:1000 dilution. An HRP-conjugated secondary antibody (from sheep, cat. #NA931V, GE Healthcare) was probed at a 1:10000 dilution. Immunoreactive bands were luminescent in a chemiluminescent substrate (Western BLoT Quant HRP substrate, cat. # T7102A, TaKaRa) and imaged using an image analyzer (LAS4000, GE Healthcare).

### Protein purification from yeast cells

Ade4-mNG, Ade4ΔIDR-mNG, and mNG were C-terminally tagged with a single Strep-tag II, expressed in budding yeast cells, and purified as follows. Yeast cells expressing the tagged protein were grown in liquid YPD medium supplemented with 0.02 mg/ml adenine sulfate at 30°C for at least 20 h. The pre-culture was diluted by 100- to 1000-fold in 100 ml of fresh YPD plus adenine medium, and cells were further grown at 30°C to 4×10^7^ cells/ml. Cells were harvested by centrifugation at 2,000×*g* and washed twice with 1 ml of extraction buffer (EB) (0.15 M NaCl, 1 mM EDTA, 1 mM dithiothreitol, and 40 mM Tris-HCl, pH 7.5) supplemented with 1 mM PMSF and cOmplete protease inhibitor cocktail (EDTA-free) (Roche, cat. # 11836170001). One gram of glass beads with a diameter of 0.5 mm (YASUI KIKAI, cat. # YGBLA05) was mixed with the cell suspension, and cells were disrupted by vigorous shaking using a beads beater (YASUI KIKAI, MB901). The crude cell extract was collected by brief centrifugation, and glass beads were washed with a small volume of EB (<0.5 ml). The crude extract and washes were combined and clarified by centrifugation twice at 20,000×*g* at 4°C for 10 min. To block endogenous biotin, the cleared lysate was supplemented with 0.2 mg/ml neutralized avidin (FUJIFILM Wako, cat. # 015-24231), incubated on ice for 30 min, and clarified by centrifugation at 20,000×*g* at 4°C for 10 min. The supernatant was loaded onto a handmade open column packed with 0.5 ml of Strep-Tactin Sepharose (IBA GmbH, cat. # 2-1201-002). After washing with 10 column volumes of EB, bound proteins were eluted with 6 column volumes of EB plus 2.5 mM D-desthiobiotin (IBA GmbH, cat. # 2-1000- 025). The protein amounts and compositions of elution fractions were analyzed by SDS- 10% PAGE and stained with FastGene Q-stain (Nippon Genetics, cat. # NE-FG-QS1). Peak fractions were pooled and concentrated using an Amicon Ultra-0.5 centrifugal filter device with a molecular weight cut-off of 30 or 10 kDa (Merck, cat. # UFC503024 or UFC501008). D-desthiobiotin in the peak fraction was diluted more than 100-fold by a repeating dilution with EB and the centrifugal concentration. The concentration of the purified protein was assessed by densitometry using bovine serum albumin (TaKaRa, cat. # T9310A) as a standard following SDS-PAGE and staining with the Q-stain. Purified Ade4-mNG and Ade4ΔIDR-mNG were supplemented with 10 mM DTT and stored on ice.

### *In vitro* condensation assay

The purified protein was diluted to the indicated concentration in assay buffer (EB plus 5 mM MgCl_2_) and supplemented with 0–15% (w/v) PEG 8000 (FUJIFILM Wako, cat. # 596-09755). In the experiments shown in Fig. S4C-E, samples were spiked with 10% (w/v) PEG of the indicated molecular size. The mixture was immediately loaded into an assay chamber composed of a glass slide and coverslip attached by double-sided tape. Images were taken after a 2-min incubation. Fluorescent condensation was imaged using the Nikon inverted fluorescent microscope (see above). Images of fluorescent condensation were analyzed by running a Fiji macro (https://github.com/masaktakaine/In-vitro-condensates-quantification). Briefly, the macro automatically detects fluorescent condensation and quantifies its area and fluorescence intensity.

### Quenching of yeast cells for a metabolomics analysis

The quenching of yeast cells before metabolite extraction followed the method reported by Kim et al. ^66^ with slight modifications. Wild-type BY4741 cells were grown to saturation in YPD medium, inoculated into synthetic medium with or without 0.02 mg/ml adenine, and then grown at 30°C for more than 16 hours. The preculture was diluted 50– 60 times with 10 ml of fresh medium, and cells were grown to approximately 1.0 × 10^7^ cells/ml at 30°C. After measuring cell density (cells/ml) with an automated cell counter (TC20, Bio-Rad), 0.9 ml of the cell culture was quenched by a rapid injection into 4.5 ml of pure methanol at −80°C. The suspension was centrifuged at 2,200×*g* at 4°C for 3 min, and the cell pellet was stored at −80°C until extraction. In samples labeled “-ade” and “+ade 24h”, cells were cultured for more than 24 h (since the pre-culture) in medium containing 0 or 0.02 mg/ml adenine, respectively. In samples labeled as “+ade 1h” and “+ade 7h”, adenine was added at a final concentration of 0.02 mg/ml 1 or 7 h before quenching, respectively.

### Metabolite extraction and widely targeted metabolomics profiling

A widely targeted metabolomic analysis was performed as previously described ^67^. In brief, each frozen sample in a 1.5-mL plastic tube was homogenized in 200 μL of 50% methanol with glass beads using a microtube homogenizer (TAITEC Corp.) at 41.6 Hz for 2 min. Homogenates were mixed with 400 μL of methanol, 100 μL of H_2_O, and 200 μL of CHCl_3_ and vortexed for 20 min. Samples were centrifuged at 20,000×*g* at 4°C for 15 min. The supernatant was mixed with 350 μL of H_2_O, vortexed for 10 min, and centrifuged at 20,000×*g* at 4°C for 15 min. The aqueous phase was collected, dried in a vacuum concentrator, and re-dissolved in 2 mM ammonium bicarbonate (pH 8.0). Chromatographic separations were performed using an Acquity UPLC H-Class System (Waters) under reverse-phase conditions with an ACQUITY UPLC HSS T3 column (100 × 2.1 mm, particle size of 1.8 μm, Waters) and under HILIC conditions using an ACQUITY UPLC BEH Amide column (100 × 2.1 mm, particle size of 1.7 μm, Waters). Ionized compounds were detected using a Xevo TQD triple quadrupole mass spectrometer coupled to an electrospray ionization source (Waters). The peak areas of target metabolites were analyzed using MassLynx 4.1 software (Waters). Metabolite signals were normalized to total ion signals of the corresponding sample. To estimate absolute PRPP concentrations, the amount of PRPP in the sample was quantified by comparisons with standard curves obtained for pure PRPP. Intracellular concentrations were then calculated using the previously reported haploid yeast cell volume (42 fL)^68^.

### MD simulation

The crystal structure of *E. coli* PPAT (PDB ID: 1ECC) was obtained from the Protein Data Bank and used for all MD simulations ^34^. All water molecules and ions were removed before simulations. To build the parameters of ligands (NH_3_, PRPP, and glutamic acid), their restrained electrostatic potential charges were calculated based on gas-phase HF/6-31(d) quantum mechanics calculations using Gaussian 16. A set of parameters were generated with a general AMBER force field using the antechamber from AMBER tools^69^.

The Amber ff14SB force field was employed for all MD simulations ^70^. Missing hydrogen atoms were added by LEaP from AMBER tools. Proteins were solvated with the TIP3P water box ^71^. The system was neutralized by replacing water with Na^+^ or Cl^-^ ions. The particle-mesh Ewald method was employed for the Coulomb interaction ^72^. Amber-formatted input files were converted for GROMACS using acpype ^73^. All MD simulations were performed using GROMACS 2019.6 ^74^. Before the production run, three-step equilibrations were performed for each system. Energy minimization was performed until 10,000 steps. This was followed by 100-ps *NVT* equilibration with the V-rescale thermostat ^75^ at 300 K under 1 atm. A 100-ps *NPT* equilibration was then conducted with Berendsen coupling ^76^ at 300 K under 1 atm. A run was performed under the *NPT* condition. The time step of each simulation was 2 fs with the SHAKE ^77^ and SETTLE ^78^ algorithms, and a snapshot was recorded every 100 ps during the production run. To examine the results of MD simulations, a Python library MDanalysis was used ^79^.

## Data analysis

Numerical data were analyzed and plotted using R studio software (R ver. 4.3.1) ^80^. Box plots show the 75th and 25th percentiles of data (interquartile range) as the upper and lower edges of the box, respectively, the median as the medial line in the box, and the 1.5×interquartile range as whiskers. The significance of differences between sets of data was indicated by asterisk(s) or *p*-values. The following abbreviations were used unless otherwise noted: *, *p* <0.05; **, *p* <0.01; ***, *p* <0.001; NS, not significant (*p* >0.05). Sparse PLS-DA was performed using the *splsda* function in the mixOmics R package^81^. A one-way ANOVA and subsequent adjustments of *p*-values by the Benjamini-Hochberg method to control the false discovery rate (FDR) were performed using Metaboanalyst 6.0 (https://www.metaboanalyst.ca/MetaboAnalyst/home.xhtml). The top 25 metabolites whose changes strongly correlated with those in the intracellular concentration of adenine were identified using the pattern hunter analysis of Metaboanalyst. All measurements were repeated at least twice to confirm reproducibility.

## Acknowledgments

The authors thank the IMCR-JURSC in Gunma University for sharing research equipment. This work was supported by research grants from the Gout and Uric Acid Foundation of Japan and by JSPS KAKENHI Grant Number JP22K06216 to M.T. This work was also supported by the joint research program of the Institute for Molecular and Cellular Regulation, Gunma University, Japan. This study used the computational resources of Cygnus provided by the Multidisciplinary Cooperative Research Program at the Center for Computational Sciences (Project Code: MOLBIO), University of Tsukuba.

## Author Contributions

MT conceived and supervised the study. MT and RM designed the experiments. MT, YY, and TN performed the experiments. RM performed MD simulations. MT, RM, YY, and TN analyzed the data. MT wrote the manuscript draft with input from RM. All authors discussed and reviewed the manuscript.

## Declaration of Interests

The authors declare no competing interests.

## Supplemental Information

### Captions for supplementary data

Data S1. Metabolomics profile of budding yeast cells grown in the absence and presence of adenine. The normalized peak area is the signal peak area corrected by the total ion chromatogram. Each group contains seven biological replicates. All data of the normalized peak area were subjected to sparse PLS-DA shown in Fig. 4B. Means and standard deviations (SD) across replicates are shown as a reference. *P*-values of the one-way ANOVA and FDR-adjusted *p*-values were calculated using MetaboAnalyst 6.0. Blue bold values indicate less than 0.05.

Data S2. Metabolic profile of 56 metabolites whose concentrations were significantly changed by adenine. Each group contains seven biological replicates. Values are the normalized peak area further converted to Z-scores across samples. Data were subjected to the heatmap analysis shown in Fig. 4C.

Data S3. Changes in purine nucleotides and their derivatives, related to Figs. 4D and S6A. Each group contains seven biological replicates. Data were converted to fold change (FC) relative to the mean at 0 h (in the absence of adenine).

### Captions for supplementary movies

**Movie S1**

Time-lapse imaging of the assembly of Ade4-mNG condensates by adenine depletion. Cells were grown in medium containing 0.02 mg/ml adenine and washed with medium lacking adenine at t = 0 min. The three cells in the upper right are shown in Fig. 1E. Scale bar = 5 µm.

**Movie S2**

Another example of cells showing the assembly of Ade4-mNG condensates by adenine depletion. Experimental conditions are the same as in Movie S1. Scale bar = 5 µm.

**Movie S3**

A control experiment of time-lapse imaging of the assembly of Ade4-mNG condensates by adenine depletion. Cells were grown in medium containing 0.02 mg/ml adenine and washed with medium containing the same concentration of adenine at t = 0 min. The rightmost cell is shown in Fig. S1B. Scale bar = 5 µm.

**Movie S4**

Time-lapse imaging of the disassembly of Ade4-mNG condensates by adenine supplementation. Cells were grown in the absence of adenine and 0.02 mg/ml adenine was supplemented at t = 0 min. The four cells in the upper left are shown in Fig. 1F. Scale bar = 5 µm.

**Movie S5**

A control experiment of time-lapse imaging of the disassembly of Ade4-mNG condensates by adenine supplementation. Cells were grown in the absence of adenine and sterile water was supplemented in place of adenine at t = 0 min. The two cells in the center are shown in Fig. S1C Scale bar = 5 µm.

**Movie S6**

Another example of cells showing Ade4-mNG condensates after a mock treatment with adenine. Experimental conditions are the same as in Movie S5. Scale bar = 5 µm.

**Movie S7**

A close examination of the motility of Ade4-mNG condensates. Another example of data shown in Fig. 1I. Cells bearing condensates were imaged on a single z-plane at 1-s intervals. The intensity of mNG is represented by an inverted grayscale. Scale bar = 5 µm.

**Movie S8**

Time-lapse video of the MD simulation of NH_3_ and PPAT molecules in the semi-holo state, related Fig. 8E. This movie was created from MD snapshots during the simulation. The state of the Arg gate is also indicated. In this example, NH_3_ was eventually released through the front channel (indicated by a circle).

**Figure S1.**
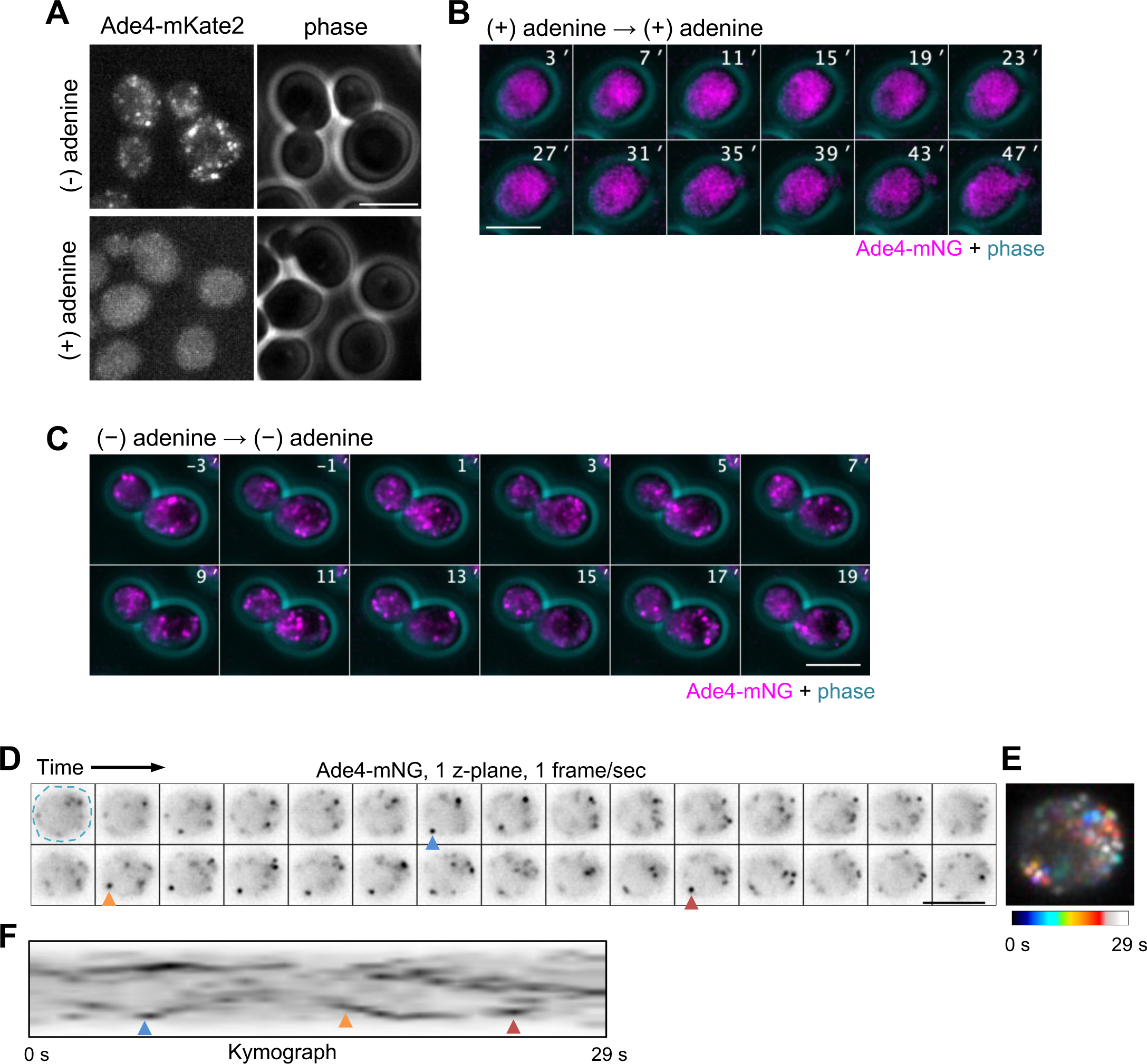
Additional data regarding the intracellular localization of Ade4. Related to Figure 1. A. Phase and RFP images of cells expressing Ade4 tagged C-terminally with the red fluorescent protein mKate2. B. A control experiment of time-lapse imaging of the assembly of Ade4-mNG particles by adenine depletion shown in Fig. 1E. Cells were grown in medium containing 0.02 mg/ml adenine and washed with medium containing the same concentration of adenine at t = 0 min. C. A control experiment of time-lapse imaging of the disassembly of Ade4-mNG particles by adenine supplementation shown in Fig. 1F. Cells were grown in the absence of adenine and sterile water was supplemented instead of adenine at t = 0 min. Scale bars = 5 µm. D-F. Another example of time-lapse imaging of Ade4-mNG particles, related to Fig. 1I- K. Experimental conditions and graph descriptions are the same as in Fig. 1I-K.

**Figure S2.**
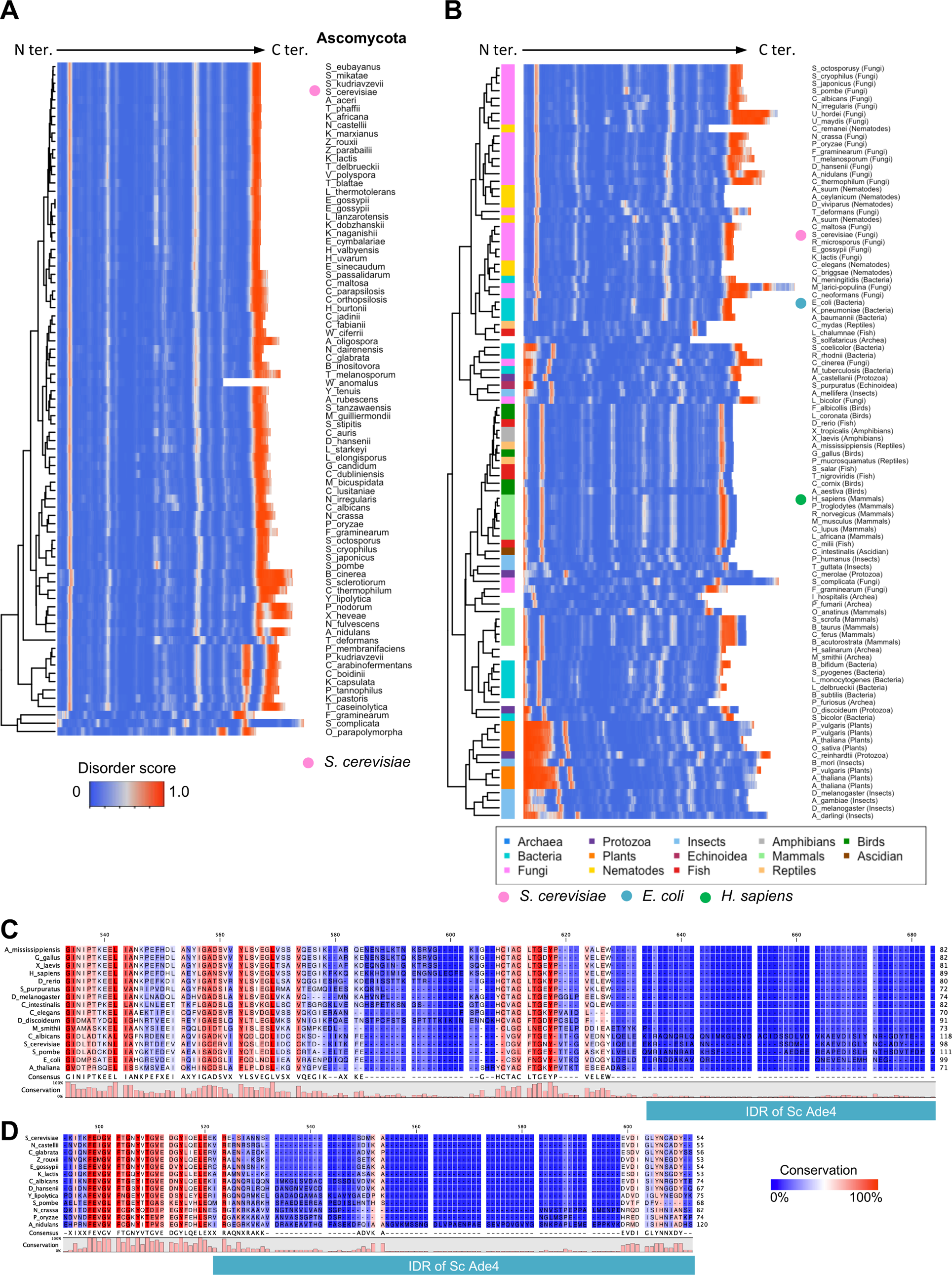
The IDR is conserved in PPATs of various species. A. The C-terminal IDR is conserved in PPATs of Ascomycota. Heatmap showing the disorder scores of PPAT amino acid sequences of 81 ascomycota. B. Heatmap showing the disorder scores of PPAT amino acid sequences of 100 organisms, including 19 ascomycota. Phylogenetic groups are color-coded on the left. C-D. Amino acid sequences of IDTs vary across species. Multiple amino acid sequence alignment of the C-terminal region of PPAT. In (C), three fungi and 13 representative organisms from the following groups are selected: archaea, bacteria, protozoa, plants, nematodes, insects, echinoidea, fish, amphibians, mammals, reptiles, birds, and ascidians. In (D), 13 species are selected from ascomycota. Green boxes indicate the region corresponding to the C-terminal IDR of Ade4.

**Figure S3.**
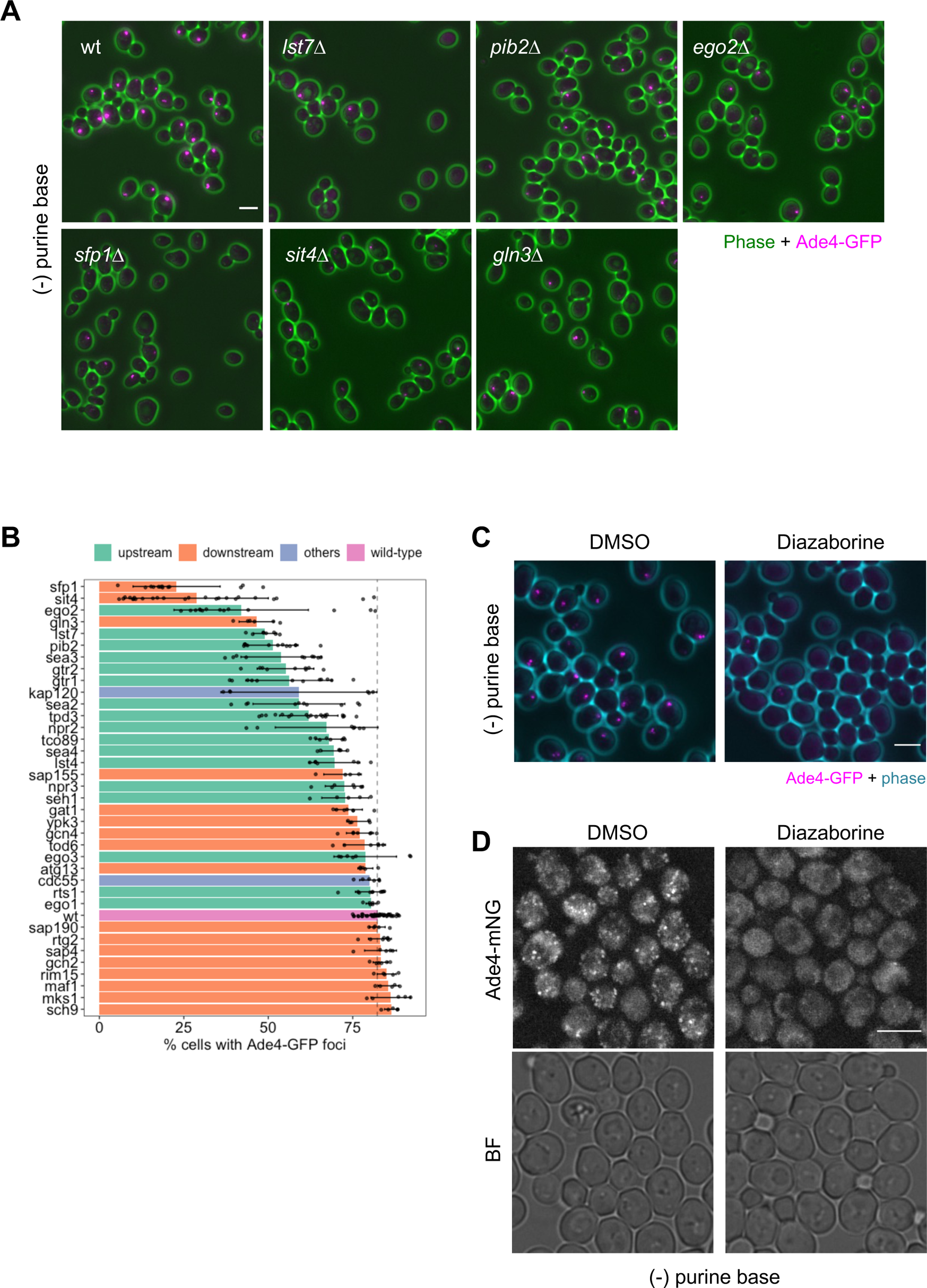
Formation of Ade4 condensates in TORC1-related mutants. Related to Figure 2. A. Assembly of Ade4-GFP foci in single gene deletion mutants of the upstream regulator and downstream effector of TORC1 signaling. Cells of the indicated genotype expressing Ade4-GFP were grown in medium containing adenine and then incubated in medium without adenine for 45 min before imaging. B. Summary of the percentage of cells showing Ade4-GFP foci in the wild-type and 37 single-gene deletion mutants. The percentage of cells with Ade4-GFP foci per field of view was plotted. Data were pooled from 2-3 independent experiments. Bars and error bars indicate the mean ± 1SD. C. The inhibition of ribosome synthesis suppressed the assembly of Ade4-GFP foci. Related to Figure 2J. D. The inhibition of ribosome synthesis suppressed the assembly of Ade4-mNG condensates. Cells expressing Ade4-mNG were grown in medium containing adenine and then incubated in medium without adenine in the presence of 0.05% (v/v) DMSO or 0.05% DMSO plus 5 µg/ml diazaborine for 45 min before imaging. Scale bars = 5 µm.

**Figure S4.**
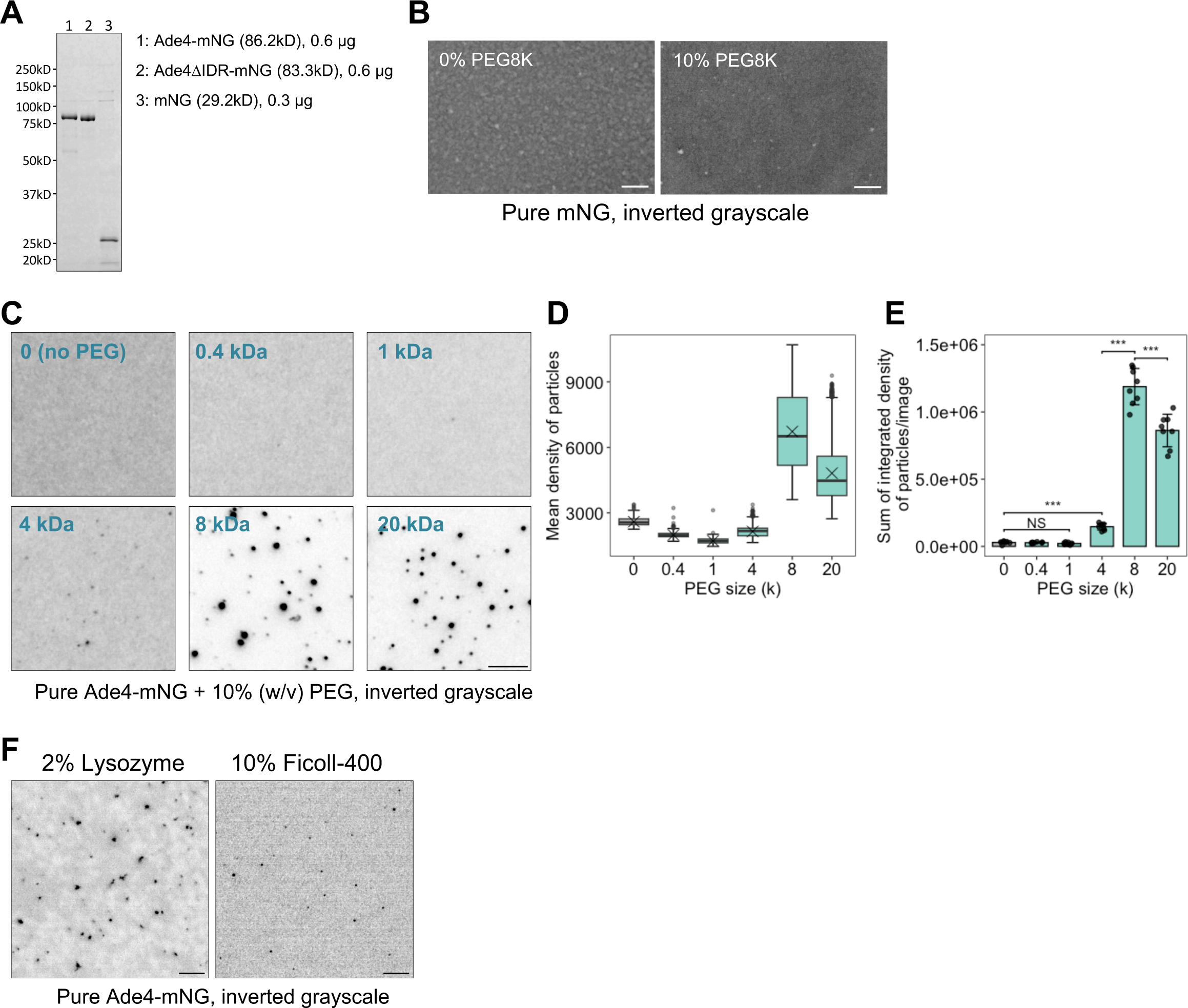
Purification and basic characterization of the Ade4 protein *in vitro*. A. Purified proteins used in the present study. Proteins were resolved by SDS-PAGE and stained with FastGene™ Q-stain. B. The purified mNG protein did not condensate into particles even under molecular crowding conditions. The supplementation of 0.14 µM mNG with 0 or 10% PEG8000 was performed. C. The *in vitro* condensation of Ade4-mNG depends on the size of PEG. The supplementation of 0.15 µM Ade4-mNG with 10% PEG of the indicated average molecular weight was performed. D-E. Quantification of data shown in (C). The mean fluorescence intensities of Ade4-mNG particles were box plotted in (D). Cross marks indicate the population mean. The integrated fluorescence intensities of particles were summed per field and plotted in (E). Bars and error bars indicate the mean ± 1SD. Seven to 8 fields of view were imaged for each condition, and an average of 399 particles per field were examined. *P*-values were calculated using the two-tailed Steel-Dwass multiple comparison test. F. Condensation of Ade4-mNG into particles with crowding agents other than PEG. The supplementation of 0.14 µM Ade4-mNG with the indicated crowding agents was performed. Scale bars = 5 µm.

**Figure S5.**
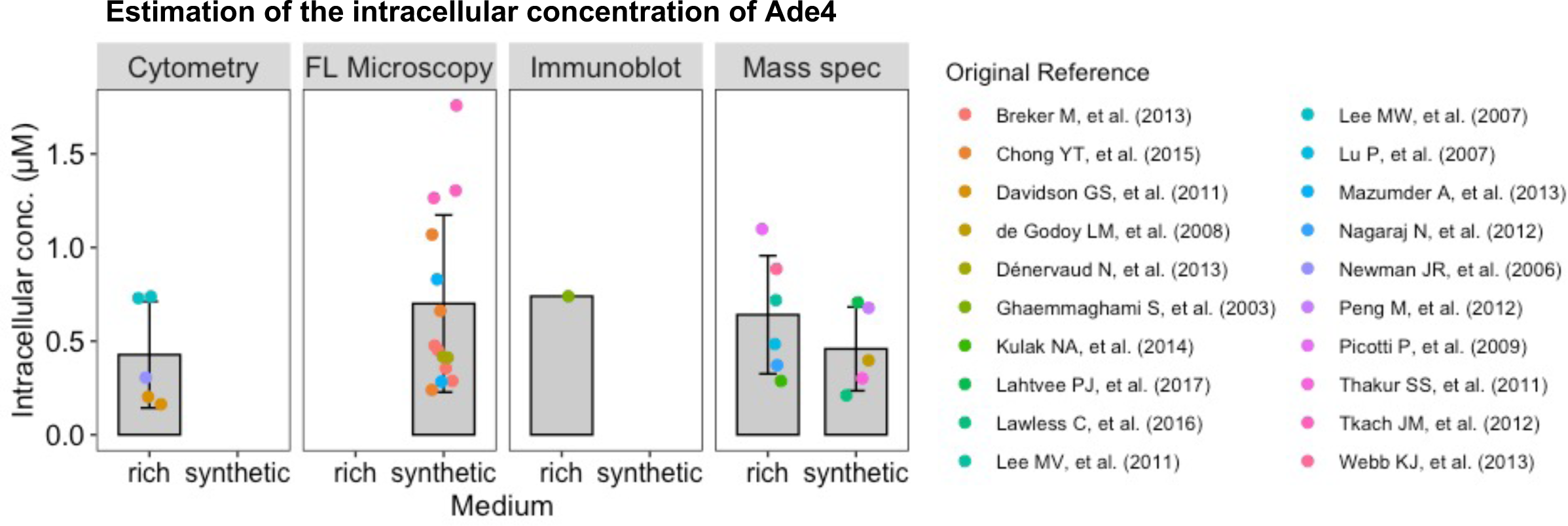
Estimation of the intracellular concentration of Ade4. Data regarding the protein abundance of the budding yeast Ade4 were downloaded from the Saccharomyces Genome Database (https://www.yeastgenome.org/locus/S000004915/protein). Data were derived from twenty quantitative genome-wide proteomic studies and normalized to a unit of molecules per cell. These values were converted to micromoles assuming that the average cytoplasmic volume of the haploid yeast cell was 42 fL^68^. Data were classified according to the measuring methods (flow cytometry, quantitative fluorescence microscopy, immunoblotting, and mass spectrometry) and the type of medium (rich or synthetic) and were then plotted. Bars and error bars indicate the mean ± 1SD.

**Figure S6.**
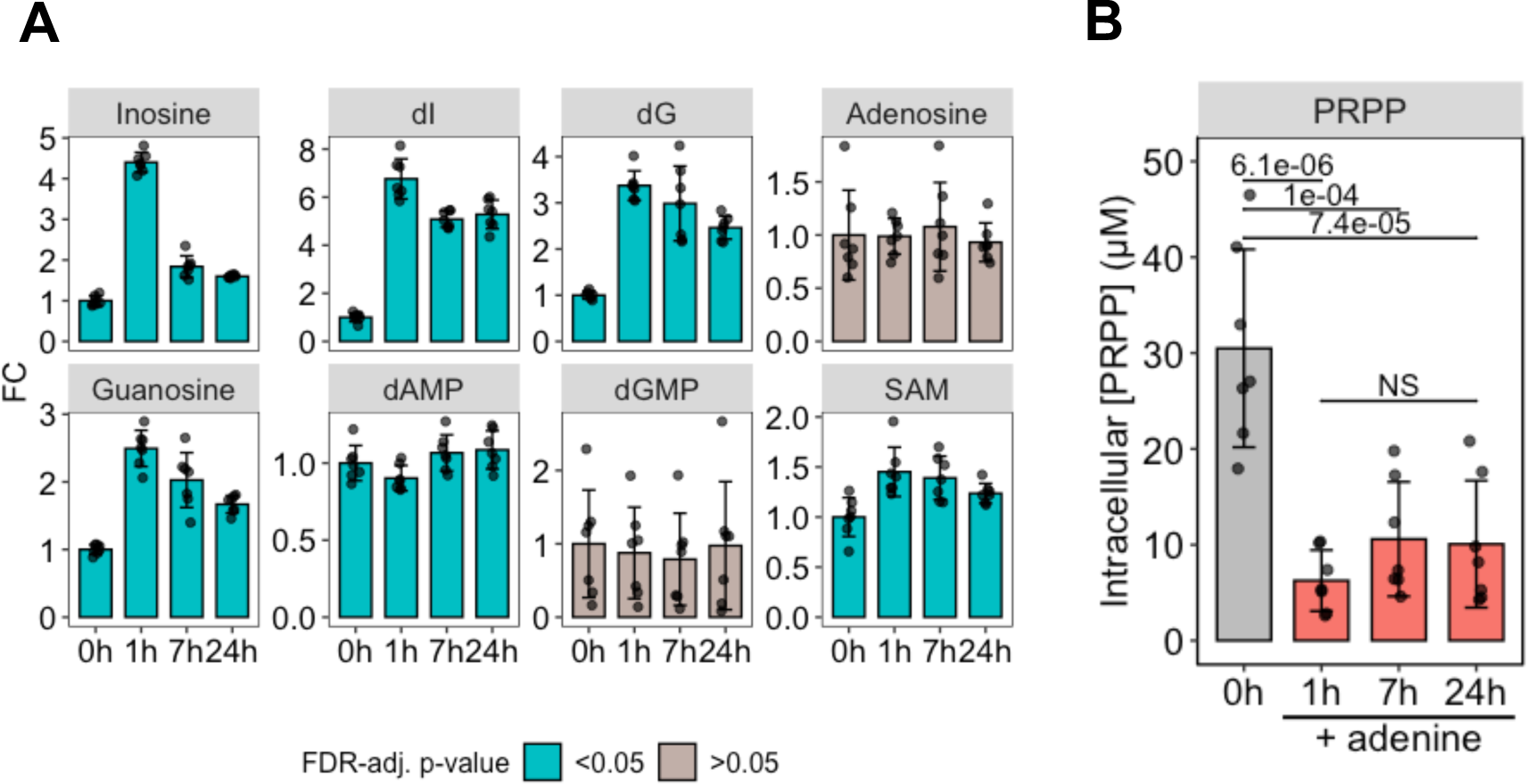
Additional data regarding the metabolomics analysis. Related to Figure 4. A. Changes in the indicated metabolites. Experimental conditions and graph descriptions are the same as in Fig. 4D. B. The intracellular concentration of PRPP decreases in the presence of environmental adenine. The amount of PRPP in each sample was converted to an intracellular concentration and plotted. *P*-values were calculated using the two-tailed Tukey-Kramer’s multiple comparison test.

**Figure S7.**
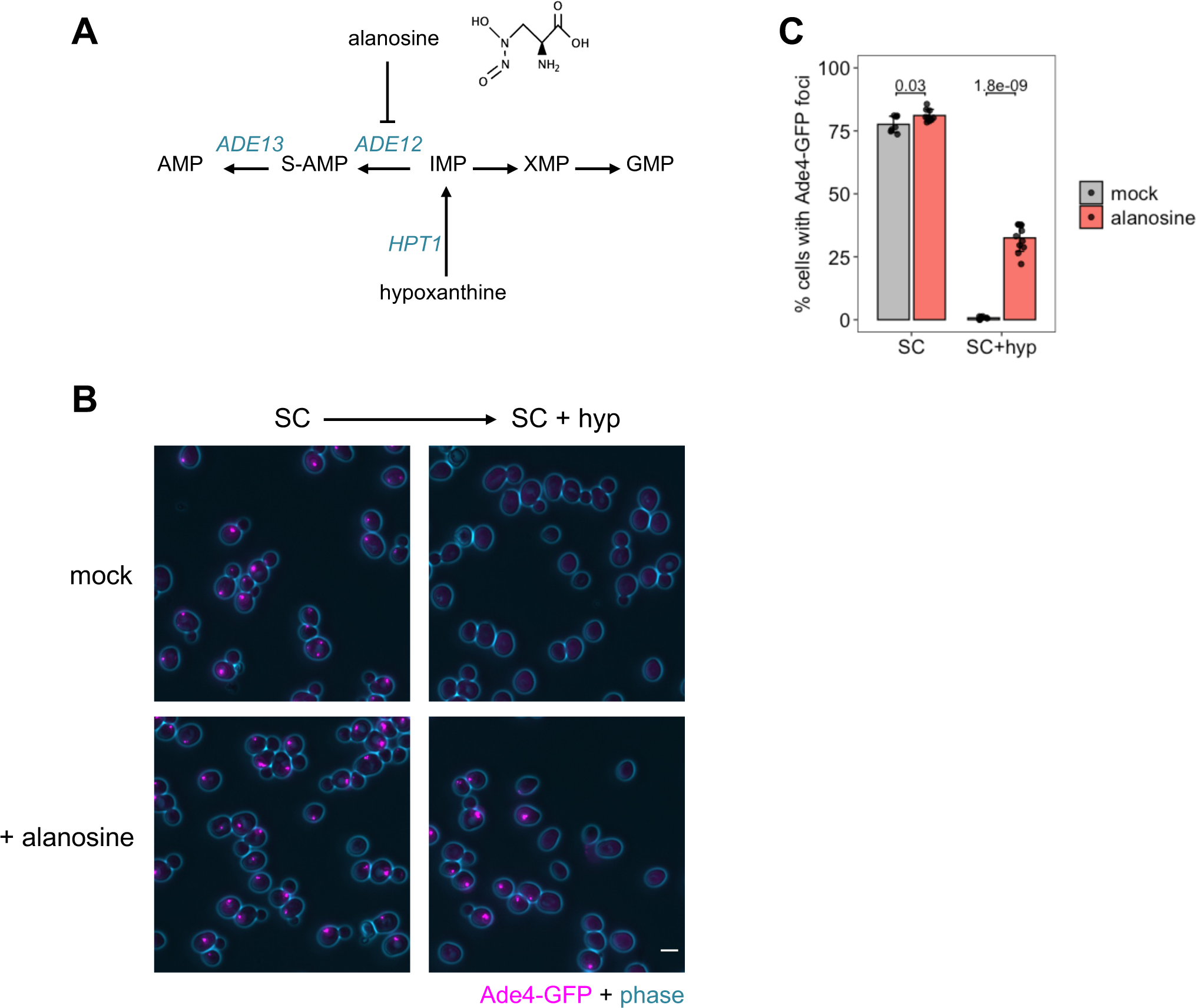
Alanosine inhibits the dissolution of Ade4 foci in the presence of hypoxanthine. Related to Figure 5. A. Alanosine inhibits the adenylosuccinate synthetase ADE12. B. Assembly of Ade4-GFP foci in the presence of alanosine. Cells expressing Ade4-GFP were incubated in medium lacking adenine in the presence of 0 (mock) or 30 µg/ml alanosine (+alanosine) for 40 min to allow for the assembly of Ade4-GFP foci and were then imaged (SC). Cells were further treated with the same medium supplemented with 24 µg/ml hypoxanthine for 15 min and then imaged again (SC + hyp). Scale bar = 5 µm. C. Quantification of data shown in (B). The percentage of cells with foci per field of view (containing 182 ± 46 cells) was quantified and plotted. Data were pooled from two independent experiments. Bars and error bars indicate the mean ± 1SD. The significance of differences was tested using the unpaired two-tailed Welch’s *t*-test. *P*-values versus the mock are shown.

**Figure S8.**
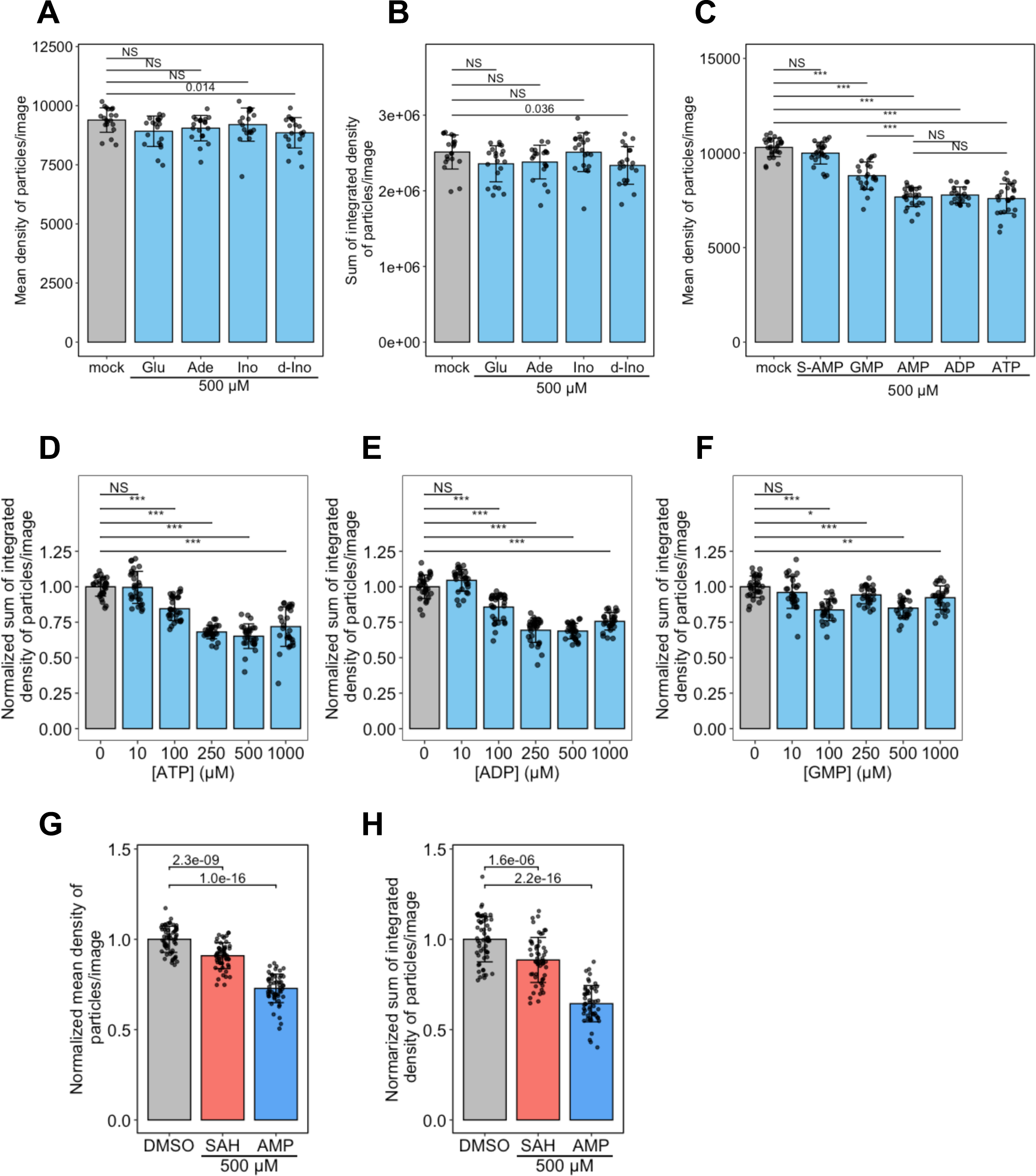
Additional data regarding the assembly of Ade4 condensates *in vitro*, related to Figure 6. A, B. Effects of purine base derivatives on the *in vitro* assembly of Ade4 condensates. The supplementation of 0.2 µM Ade4-mNG with 10% PEG8000 was performed in the presence of the indicated compound. Glucose (Glu) was used as a negative control. The integrated fluorescence intensities of condensates were summed per field and plotted in (A). The mean fluorescence intensities of Ade4-mNG condensates were averaged per field and plotted in (B). Bars and error bars indicate the mean ± 1SD. Twenty fields of view were imaged for each condition from two independent preparations, and an average of 1452 particles per field were examined. *P*-values were calculated using the two-tailed Steel’s multiple comparison test. Glu, glucose; Ade, adenine; Ino, inosine; d-Ino, deoxy-inosine. C. Effects of purine nucleotides on the *in vitro* assembly of Ade4 condensates. Data were from the same experiment shown in Figure 6A. The mean fluorescence intensities of Ade4-mNG condensates were averaged per field and plotted. Bars and error bars indicate the mean ± 1SD. *P*-values were calculated using the two-tailed Tukey-Kramer’s multiple comparison test. D-F. The supplementation of 0.2 µM Ade4-mNG with 10% PEG8000 was performed in the presence of the indicated concentrations of ADP (D), ATP (E), and GMP (F). The integrated fluorescence intensities of the condensates were summed per field, normalized to the mean at 0 µM, and plotted. Bars and error bars indicate the mean ± 1SD. From two independent preparations, 25-31 fields of view were imaged for each condition, and an average of 1200 particles per field were examined. *P*-values were calculated using the two-tailed Steel’s multiple comparison test. G-H. Inhibitory effect of SAH on the *in vitro* condensation of Ade4-mNG. The supplementation of 0.2 µM Ade4-mNG with 10% PEG8000 was performed in the presence of 0.5% (v/v) DMSO and 500 µM SAH or AMP and imaged. The average fluorescence intensities (G) and integrated fluorescence intensities (H) of the particles summed per field were normalized to the mean of the DMSO alone and plotted. Bars and error bars indicate the mean ± 1SD. From 4 independent preparations, 53-57 fields of view were imaged for each condition. The *p*-values were calculated using the two-tailed Dunnett’s multiple comparison test.

**Figure S9.**
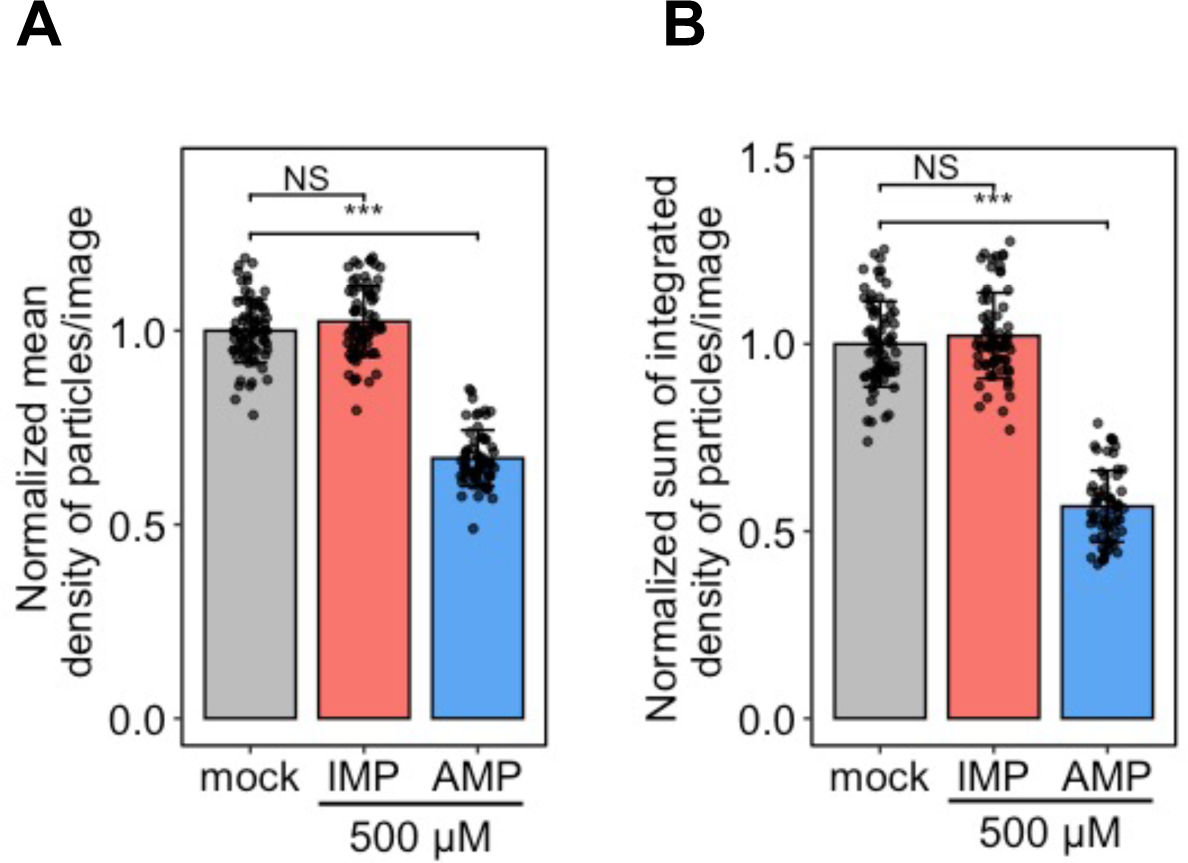
Effects of IMP on the assembly of Ade4 condensates *in vitro*. A-B. IMP did not affect the *in vitro* condensation of Ade4-mNG. The supplementation of 0.2 µM Ade4-mNG with 10% PEG8000 was performed in the presence of 500 µM IMP or AMP. The average fluorescence intensities (A) and integrated fluorescence intensities (B) of the condensates summed per field were normalized to the mean of the mock, and plotted. Bars and error bars indicate the mean ± 1SD. From 4 independent preparations, 62-72 fields of view were imaged for each condition. *P*-values were calculated using the two-tailed Steel’s multiple comparison test.

**Figure S10.**
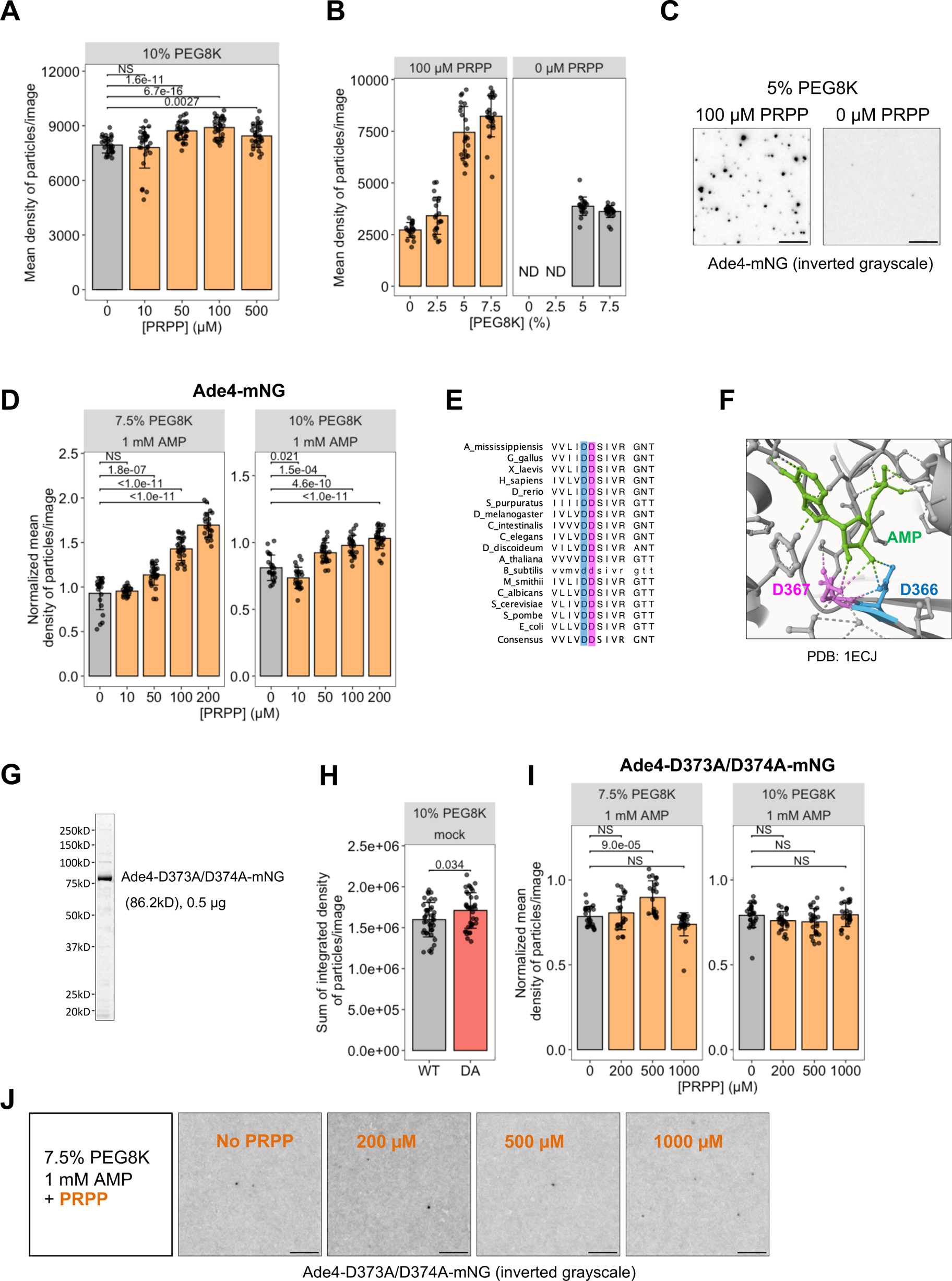
PRPP facilitates the assembly of Ade4 condensates, related to Figure 7. A. PRPP augments the *in vitro* assembly of Ade4 condensates. Data were from the same experiment shown in Figure 7A. The mean fluorescence intensities of Ade4-mNG condensates were averaged per field and plotted. Throughout the figure, bars and error bars indicate the mean ± 1SD. *P*-values were calculated using the two-tailed Steel’s multiple comparison test. B, C. PRPP promotes the formation of Ade4 condensates *in vitro* at suboptimal PEG concentrations. Data were from the same experiment shown in Figure 7B, C. The mean fluorescence intensities of Ade4-mNG condensates were averaged per field and plotted in (B). ND, not determined. Representative images at 5% PEG are shown in (C). D. PRPP antagonizes the inhibitory effects of AMP. Data were from the same experiment shown in Figure 7D-E. The mean fluorescence intensities of Ade4-mNG condensates were averaged per field, normalized to the mean of the mock control (no AMP and no PRPP), and plotted. *P*-values were calculated using the two-tailed Steel’s multiple comparison test. E. The amino acid residues of the PRPP binding motif in PPAT are well conserved among species. F. Interactions between an AMP and specific amino acid residues in the PRPP binding motif in the *E. coli PPAT* (pdb file: 1ecj). The dotted line indicates a hydrogen bond. G. The purified Ade4-D373A/D374A-mNG (Ade4-DA-mNG) protein. Proteins were resolved by SDS-PAGE and stained with FastGene™ Q-stain. H. Ade4-DA-mNG condensated into particles under molecular crowding conditions. The supplementation of 0.2 µM Ade4-mNG (WT) and Ade4-DA-mNG (DA) with 10% PEG8000 was performed and imaged. The integrated fluorescence intensities of the condensates were summed per field and plotted. From 2 independent preparations, >36 fields of view were imaged for each condition. *P*-values were calculated using the two-tailed Mann-Whitney U test. I, J. PRPP did not antagonize AMP in the condensate formation of the DA mutant. Data were from the same experiment shown in Figure 7K. The mean fluorescence intensities of Ade4-D373A/D374A-mNG condensates were averaged per field, normalized to the mean of the mock control (no AMP or PRPP), and plotted in (I). *P*-values were calculated using the two-tailed Steel’s multiple comparison test. Representative images at 7.5% PEG are shown in (J). Scale bars = 5 µm.

**Figure S11.**
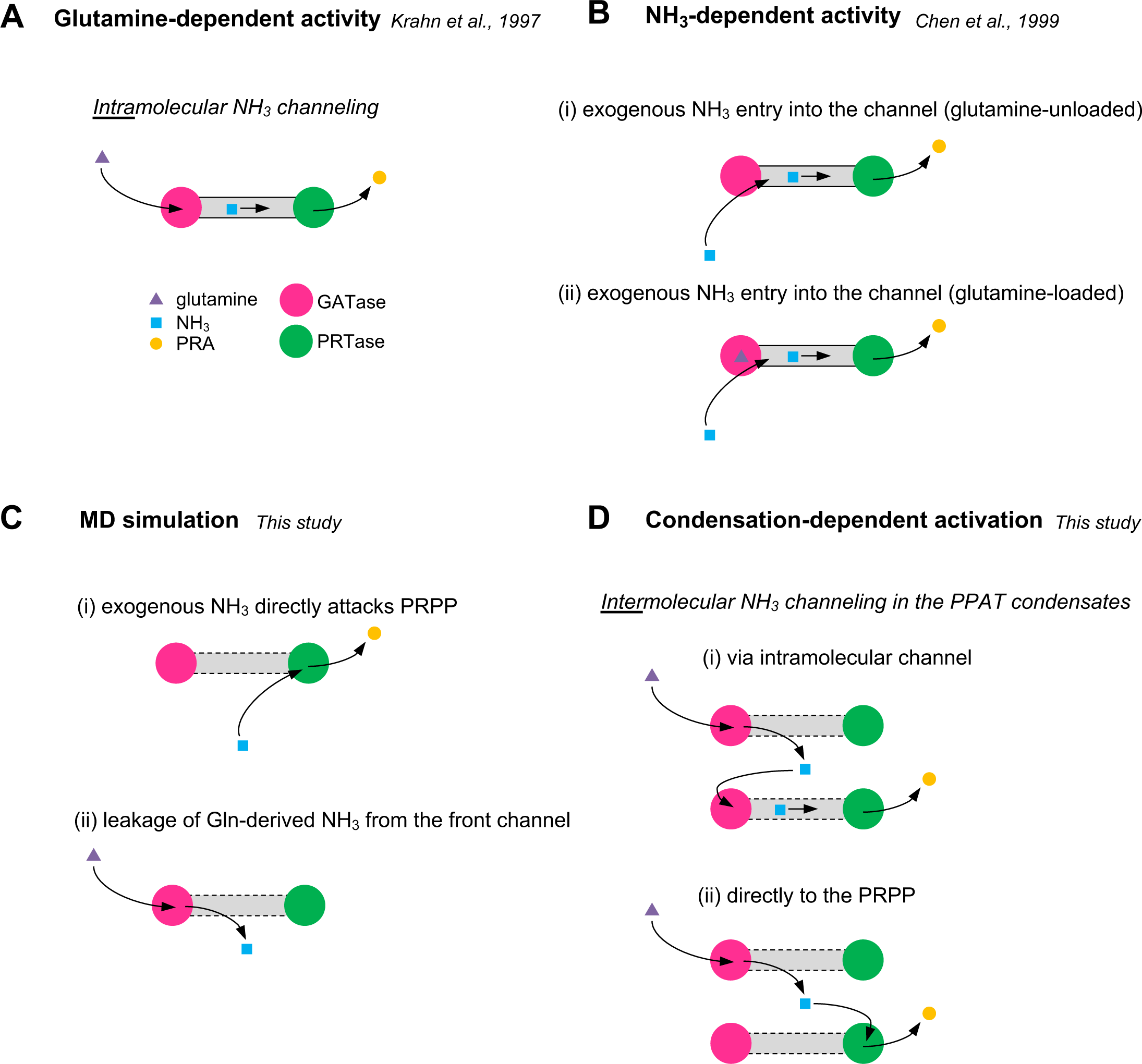
Possible enzymatic reaction pathways of PPAT. A. Intramolecular channeling of glutamine-derived NH_3_ proposed by Krahn et al.^34^. B. Mechanism of NH_3_-dependent activity proposed by Chen et al.^53^. External NH_3_ enters the channel from the glutamine site irrespective of glutamine binding. C. The MD simulation in the present study suggests that glutamine-derived NH_3_ leaks through the front channel following the attack of exogenous NH_3_. D. Condensation-dependent activation of Ade4 proposed by the present study. Glutamine-derived NH_3_ leaked from the front channel rapidly diffuses away. In the Ade4 condensate, the opportunity for NH_3_ to be captured by other molecules and further reacted with PRPP will increase. This activation may be described as intermolecular NH_3_ channeling. PRPP is not depicted in the schema.

## Notes

### Competing Interest Statement

The authors have declared no competing interest.

### Summary of Updates

Minor corrections to text and graphs.

